# The nucleus is a quality control center for non-imported mitochondrial proteins

**DOI:** 10.1101/2020.06.26.173781

**Authors:** Viplendra P.S. Shakya, William A. Barbeau, Tianyao Xiao, Christina S. Knutson, Adam L. Hughes

## Abstract

Mitochondrial import deficiency causes cellular stress due to the accumulation of non-imported mitochondrial precursor proteins. Despite the burden mis-localized mitochondrial precursors place on cells, our understanding of the systems that dispose of these proteins is incomplete. Here, we catalog the location and steady-state abundance of mitochondrial precursor proteins during mitochondrial impairment in *S. cerevisiae*. We find that a number of non-imported mitochondrial proteins localize to the nucleus, where they are eliminated by proteasome-based nuclear protein quality control. Recognition of mitochondrial precursors by the nuclear quality control machinery requires the presence of an N-terminal mitochondrial targeting sequence (MTS), and impaired breakdown of precursors leads to their buildup in nuclear-associated foci. These results identify the nucleus as a key destination for the disposal of non-imported mitochondrial precursors.

## Main Text

Mitochondrial dysfunction is a hallmark of aging and associated with many age-related and metabolic diseases (*1*). Mitochondrial impairment disrupts metabolic pathways housed within the mitochondrion, and also prevents the import of thousands of mitochondrial resident proteins that rely on an efficient mitochondrial membrane potential for translocation into the organelle (*2–4*). Recent studies have shown that non-imported mitochondrial precursor proteins are toxic for cells, and identified several cellular pathways that combat this stress by disposing or triaging non-imported precursors (*5–11*). However, despite these recent advances, only a fraction of the non-imported mitochondrial proteome has been analyzed under conditions of mitochondrial impairment. Thus, our understanding of the fate of non-imported mitochondrial precursors remains incomplete. Here, using microscopy and immunoblot-based screens in *S*. *cerevisiae*, we show that non-imported mitochondrial proteins accumulate in many regions of the cell upon mitochondrial depolarization, and identify the nucleus as an important quality control destination for non-imported mitochondrial precursors. We find that many mitochondrial proteins localize to the nucleus upon import failure, where they are subjected to proteasome-dependent destruction via redundant action of the E3 ubiquitin ligases San1, Ubr1, and Doa10. When degradation capacity is exceeded, mitochondrial precursors are sequestered into nuclear-associated protein aggregates. We show that the N-terminal mitochondrial targeting sequence (MTS) (*12*) is necessary for non-imported precursor protein induced-toxicity, degradation, and sequestration into aggregates, but dispensable for nuclear localization. The presence of an MTS is also required for degradation and toxicity of non-imported proteins that localize to cellular regions other than the nucleus, implicating the MTS as a major driver of non-imported precursor toxicity. Finally, we show that nuclear accumulation of non-imported precursors arises during cellular aging. Overall, this work demonstrates that non-imported mitochondrial proteins exhibit numerous fates within cells, and identifies the nucleus as an important quality control destination for non-imported mitochondrial precursor proteins.

We previously showed that the mitochondrial network undergoes extensive fragmentation and depolarization during replicative aging in budding yeast, which is defined as the number of times an individual yeast cell undergoes division (*13*). In our earlier work, we utilized an endogenously tagged version of the mitochondrial outer membrane (OM) protein Tom70-GFP to visualize the mitochondrial network. In contrast to Tom70, which does not rely on mitochondrial membrane potential for its mitochondrial localization (*14*), functional, endogenously GFP-tagged Ilv2 (fig. S1A), a key mitochondrial matrix enzyme in isoleucine and valine biosynthesis (*15*), exhibited dual localization in replicatively aged yeast cells. In addition to a pattern consistent with mitochondrial tubules, Ilv2-GFP also localized to the nucleus in over 80% of aged cells, as indicated by diffuse GFP fluorescence within a region surrounded by the nuclear pore protein Nup49-mCherry (Fig. 1A).

**Fig. 1.**
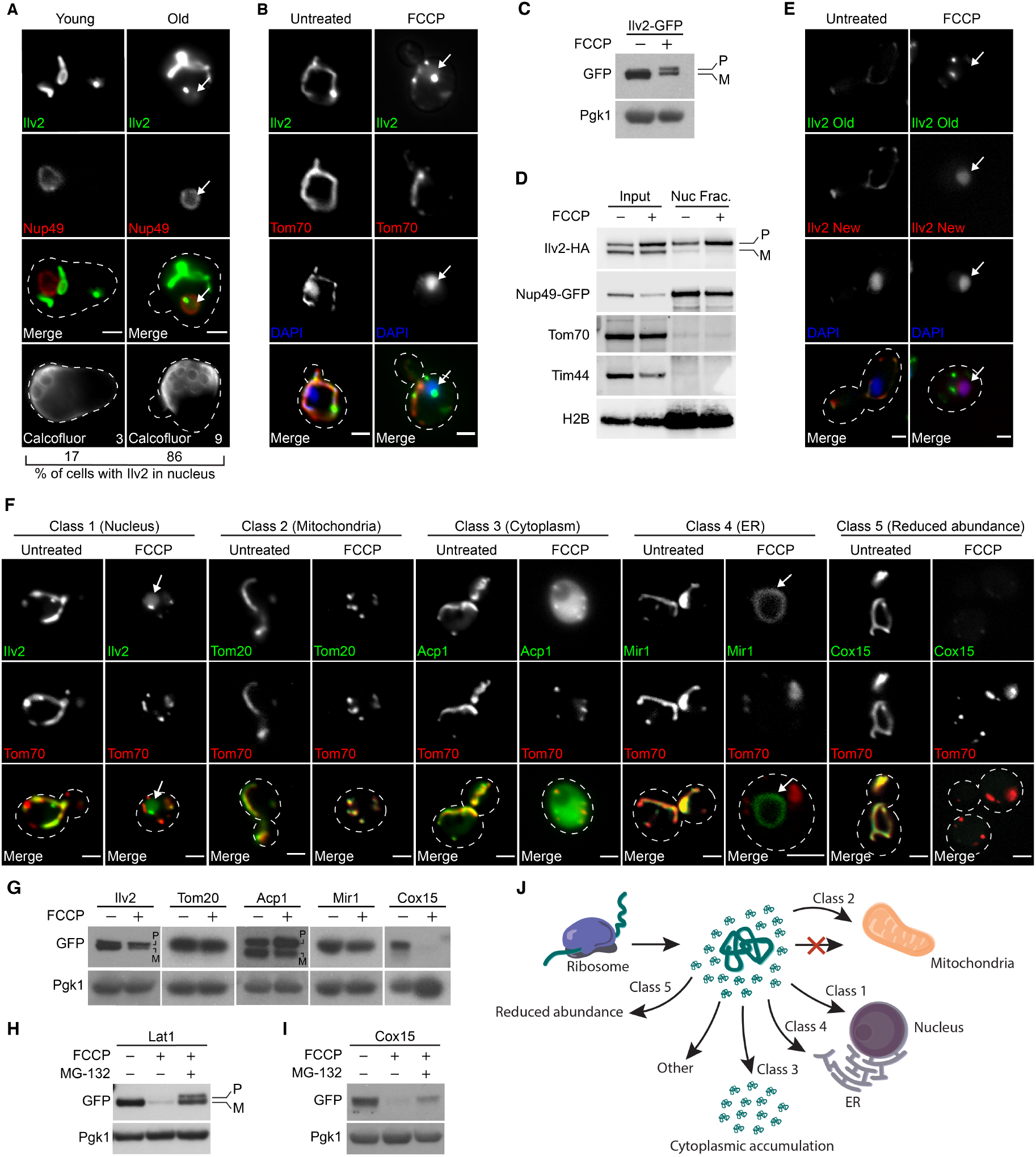
The nucleus in one of several fates for non-imported mitochondrial proteins. (**A**) Representative images of old and young yeast expressing the indicated Ilv2-GFP and nuclear marker Nup49-mCherry. Percentage of cells with Ilv2 in the nucleus (n = 30 cells) and age of representative cell (determined by bud scar counting) are indicated in bottom panels. Bud scars stained with calcofluor. (**B)** Yeast expressing Ilv2-GFP and Tom70-mCherry −/+ FCCP. (**C**) Western blots of yeast expressing Ilv2-GFP −/+ FCCP. P = precursor, M = mature in all instances. Pgk1 = loading control in all instances. (**D**) Western blot showing enrichment of the precursor form of Ilv2-HA in the nuclear fraction. Nup49-GFP and H2B = nuclear markers, Tom70 and Tim44 = mitochondrial markers. (**E)** RITE-tagged cells treated with β-estradiol at time of FCCP addition to initiate Cre/lox switching of Ilv2 epitope tag from GFP (old) to RFP (new). (**B** and **E**) Nucleus stained with DAPI. (**F)**, Yeast expressing the indicated mCherry or GFP-tagged mitochondrial proteins −/+ FCCP. (**G** to **I**) Western blots of yeast expressing the indicated GFP-tagged mitochondrial proteins −/+ FCCP (G) or −/+ FCCP −/+ MG-132 (H, I). (**J**) Summary of non-imported mitochondrial protein fates. All scales bars = 2μm. Arrows denote nucleus (**A**, **B**, **E**, and **F**, class 1) or ER (**F**, class 4).

Mitochondrial depolarization is a hallmark feature of aged yeast (*13*). Because depolarization is prominent, and the import of Ilv2 requires a mitochondrial inner membrane (IM) potential, we wondered whether the fraction of Ilv2-GFP localized to the nucleus represented a non-imported precursor pool of this protein. Consistent with that idea, treatment of cells with the mitochondrial IM depolarizing agent trifluoromethoxy carbonyl cyanide phenylhydrazone (FCCP) (*16*) also caused Ilv2-GFP accumulation in the nucleus, which was marked with 4’,6-Diamidine-2’-phenylindole dihydrochloride (DAPI) (Fig. 1B). Nuclear localization did not result from GFP-tagging, as indirect immunofluorescence showed a similar nuclear localization of C-terminally FLAG-tagged Ilv2 in FCCP treated cells (fig. S1B). Furthermore, Ilv2-GFP localized to the nucleus in cells conditionally depleted of the essential OM protein import channel Tom40 (*17, 18*) (fig. S1C and D), indicating that nuclear localization was not caused by off-target effects of FCCP, but was specific to defects in mitochondrial protein import.

We hypothesized that the nuclear pool of Ilv2 likely represented a fraction of the protein that failed to import into mitochondria. Consistent with that idea, western blot analysis revealed that a higher-molecular weight form of Ilv2-GFP and Ilv2-HA accumulated in cells treated with FCCP (Fig. 1C and fig. S1E). Mitochondrial proteins such as Ilv2 are synthesized with an N-terminal MTS extension that is proteolytically removed from the mature peptide only after they transit the mitochondrial IM (*19, 20*). Thus, the higher-molecular weight form of Ilv2 in FCCP treated cells likely represents the immature, precursor form of the protein. In support of the idea that the non-imported pool of Ilv2 localizes to the nucleus, the precursor form of Ilv2-HA was specifically enriched in nuclear fractions isolated from FCCP-treated cells, while other mitochondrial proteins, including Tom70, Tim44, as well as the mature form of Ilv2, were excluded (Fig. 1D). Additionally, we utilized the Recombination-Induced Tag Exchange (RITE) system (*21*) to examine the fate of both old and newly synthesized Ilv2 in the same cell, and found that only newly synthesized Ilv2 localized to the nucleus upon FCCP treatment, while Ilv2 already present in mitochondria did not (Fig. 1E). Collectively, these results indicate that when the translocation of Ilv2 into mitochondria is blocked by genetic or pharmacologic impairment of mitochondrial import, the non-imported precursor form of Ilv2 alternatively localizes to the nucleus.

We next sought to determine the extent to which non-imported proteins localize to the nucleus in cells lacking efficient mitochondrial import. To address this question in a systematic manner, we imaged a collection of yeast strains expressing 526 distinct mitochondrial proteins with carboxy-terminal GFP fusions from their endogenous loci in the absence or presence of FCCP. These strains were derived from the yeast GFP collection (*22*) and co-expressed Tom70-mCherry, a mitochondrial OM marker that localizes to mitochondria independently of the membrane potential (*14, 23*). We found that 6.3% of the mitochondrial proteins analyzed behaved like Ilv2, exhibiting nuclear localization in FCCP treated cells (class 1, Fig. 1F and table S1). Additionally, we identified four other major outcomes for mitochondrial proteins after membrane depolarization (Fig. 1F and table S1). These included continued localization to the mitochondrion (class 2, e.g., Tom20, 8.4% of all proteins), accumulation in the cytoplasm (class 3, e.g., Acp1, 36.1% of all proteins), localization to the endoplasmic reticulum (ER) (class 4, e.g., Mir1, 2.9% of all proteins), and reduced overall abundance to the point of being nearly undetectable (class 5, e.g., Cox15, 42.0% of all proteins). A subset of proteins (4.3%) localized to regions of the cell distinct from these five major classes upon FCCP treatment and associated with unidentified cellular membranes and foci (table S1). We validated representatives from each class and confirmed ER localization of class 4 proteins via co-localization with the ER marker Sec61-mCherry (Fig. 1F fig. S1F). As with Ilv2-GFP, identical fates occurred for all classes of proteins in cells conditionally depleted of the essential OM protein import channel Tom40 (*17, 18*) (fig. S1G), as well as in cells expressing FLAG-tagged versions of the proteins (fig. S1H), indicating the observed changes were not caused by off target effects of FCCP or the presence of a GFP tag.

We concurrently analyzed steady-state protein abundance via western blotting of the same set of GFP-tagged mitochondrial proteins in the absence and presence of FCCP, as this approach provided useful information about the state of Ilv2 in the nucleus. In general, steady-state levels of proteins localized to the mitochondrion, cytoplasm, and ER were unchanged or partially reduced with FCCP (Fig. 1G, table S1). Proteins that localized to the nucleus or became undetectable often either moderately or strongly decreased in abundance upon FCCP treatment, respectively (Fig. 1G, table S1). The decline in class 5 protein abundance was either completely or partially blunted by proteasome inhibition via MG-132 depending on the individual protein substrate (Fig. 1, H and I), implicating the proteasome in their destruction. Furthermore, as with Ilv2, precursor forms of Acp1 and Lat1 (class 3 and 5) were visible in the presence of FCCP (Fig. 1, G and H), and C-terminally HA-tagged versions of representatives from each of the five classes showed identical alterations in protein levels as the GFP-tagged versions (fig. S1, I to K).

Overall, our screen revealed several patterns amongst the proteins that comprised each screen class, and many of our observations aligned well with those from previous studies (Fig. 1J). Nuclear-localized class 1 proteins were predominantly mitochondrial matrix enzymes, including numerous members of the TCA cycle. Most class 2 proteins that continued to localize to depolarized mitochondria were mitochondrial OM proteins that do not require a membrane potential for mitochondrial targeting (*4*). Class 3 (cytoplasm) proteins were largely soluble proteins, several of which (e.g., Idh1, Idh2, Mss116, and Cis1) were previously found to be enriched in cytosolic extracts isolated from mitochondrial import-deficient yeast (*5, 7*). ER-localized class 4 proteins were generally integral IM and OM proteins, some of which were previously reported to localize to the ER in cells with compromise mitochondrial import(*9*). Finally, class 5, the largest of the classes, consisted of both soluble and membrane-bound mitochondrial proteins.

We next wanted to understand the basis for the nuclear localization of non-imported mitochondrial proteins in the absence of functional mitochondrial import. The eukaryotic nucleus harbors a large proportion of cellular proteasomes, and is a quality control destination for misfolded proteins (*24*). Because the overall abundance of nuclear-localized mitochondrial proteins declined during FCCP treatment, we tested whether non-imported mitochondrial precursor proteins were directed to the nucleus for proteasomal degradation. In support of that idea, the decline in steady-state levels of Ilv2-GFP and Ilv2-HA upon FCCP treatment was blunted in the presence of proteasome inhibitor MG-132 (Fig. 2A and fig. S2A). Ilv2 decline was also prevented in strains lacking a combination of three E3 ubiquitin ligases that operate in nuclear-associated protein quality control, San1 (*25*), Ubr1 (*26*), and Doa10 (*27, 28*) (E3 KO) (Fig. 2B and fig. S2B). No combination of single or double knockouts completely prevented loss of Ilv2 upon mitochondrial depolarization, suggesting these ligases act redundantly to promote non-imported mitochondrial protein clearance (fig. 2C). Importantly, the addition of proteasome inhibitor or deletion of the aforementioned E3 ligases each led to a marked elevation in the higher molecular weight precursor form of Ilv2 in the presence of FCCP, suggesting the immature, Ilv2 precursor was the form of the protein specifically marked for proteasome clearance (Fig. 2, A and B, fig. S2, A and B). In line with this observation, cycloheximide-chase analysis demonstrated that the half-life of the Ilv2 precursor form was altered in the E3 KO strain, while the mature form was unaffected (Fig. 2C and fig. S2D). Furthermore, ubiquitin immunoprecipitation assays indicated that Ilv2 was ubiquitylated in the presence of FCCP in a San1, Ubr1, and Doa10-dependent manner (Fig. 2D). Proteasome-dependent degradation of a non-nuclear class 5 substrate (Lat1) was unaffected in the E3 KO strain, indicating that additional E3 ligases promote clearance of non-nuclear localized mitochondrial precursors (fig. S2E). Finally, we found that our observations extend beyond Ilv2, as two other nuclear candidates identified in our screen were also eliminated in a proteasome and San1/Ubr1/Doa10-dependent manner (fig. S2, F to K). Thus, a subset of non-imported mitochondrial proteins are subjected to nuclear-associated protein quality control when their import into mitochondria is impaired.

**Fig. 2.**
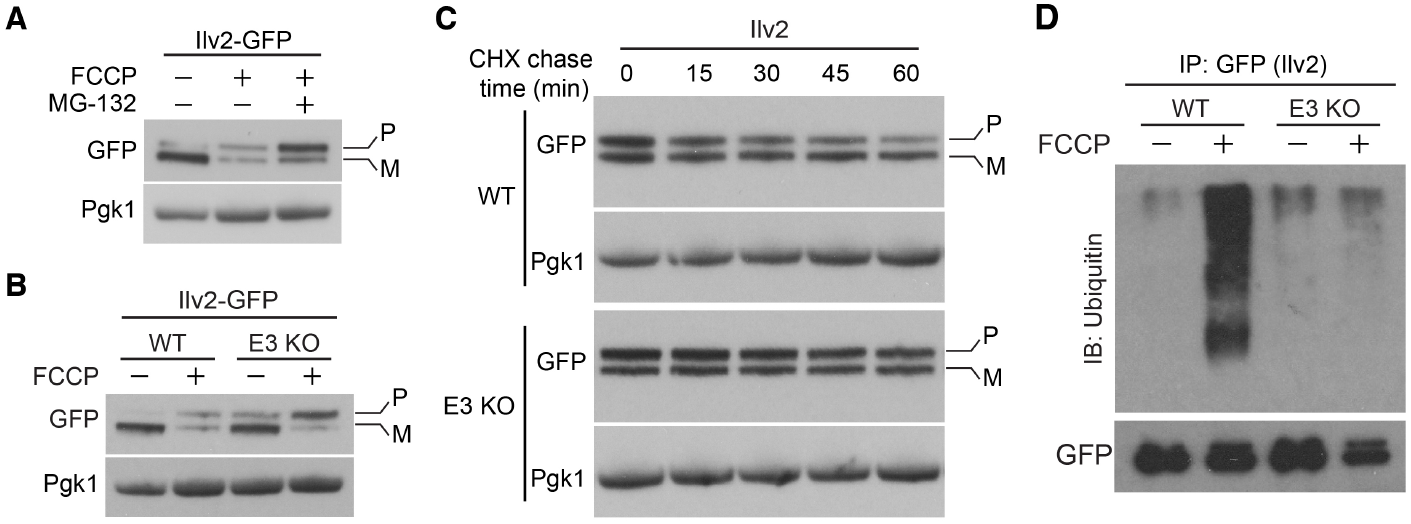
Nuclear protein quality control clears unimported mitochondrial proteins. (**A**) Western blot of yeast expressing Ilv2-GFP −/+ FCCP −/+ MG-132. (**B**) Western blot of yeast expressing Ilv2-GFP −/+ FCCP in wild-type (WT) and E3 KO strains. (**C**) Western blots showing cycloheximide (CHX) chase of Ilv2-GFP in WT and E3 KO strains in the presence of FCCP. (**D**) Western blot showing ubiquitylation of immunoprecipitated Ilv2-GFP −/+ FCCP in WT and E3 KO strains. Pgk1 = loading control. E3 KO = *san1Δ ubr1Δ doa10Δ*. P = precursor and M = mature in all instances.

As the toxicity of non-imported precursor proteins is now well documented (*5*), we wondered whether failure to destroy nuclear-localized non-imported precursors would compromise cellular health. To test this idea, we compared the growth of wild type and the aforementioned E3 KO strains in the absence and presence of FCCP. We observed no growth defect in single, double, or triple E3 ligase knockout strains (Fig. 3A, fig. S3A), suggesting redundant systems may act to mitigate the toxicity of nuclear-localized non-imported proteins. Consistent with that idea, we noticed that in addition to diffuse nuclear localization, a portion of Ilv2-GFP accumulated in nuclear-associated foci that resembled previously described juxtanuclear (JUNQ) (*29*) or intranuclear (INQ) (*30*) protein aggregate compartments (Fig. 3B). These foci were adjacent to the DAPI-stained nucleus, excluded the mitochondrial marker Tom70-mCherry, and were present in a high percentage of FCCP-treated cells (Fig. 3C). Prior studies showed that misfolded proteins can be sequestered into nuclear associated aggregates when their proteasomal clearance is impaired (*29, 30*). Consistent with that idea, the intensity of Ilv2-GFP foci increased in the E3 KO strain (Fig. 3D). Moreover, Dld1 and Dld2, which are degraded more robustly than Ilv2, also localized to nuclear-associated protein aggregates, but only in strains lacking the E3 ligase degradation machinery (fig. S3, B to E). We were unable to identify a mutation that blocked localization to these puncta. However, we did find that a two-fold increase in expression of Ilv2-GFP from a single copy plasmid resulted in constitutive localization of Ilv2-GFP to the nucleus and nuclear associated protein foci (see Fig. 4, B to D), and resulted in severe growth defects in both wild-type and E3 KO strains (Fig. 3E). These results indicate that non-imported nuclear-localized mitochondrial proteins are toxic, and that proteasome destruction and aggregate sequestration may act in coordination to mitigate this toxicity.

**Fig. 3.**
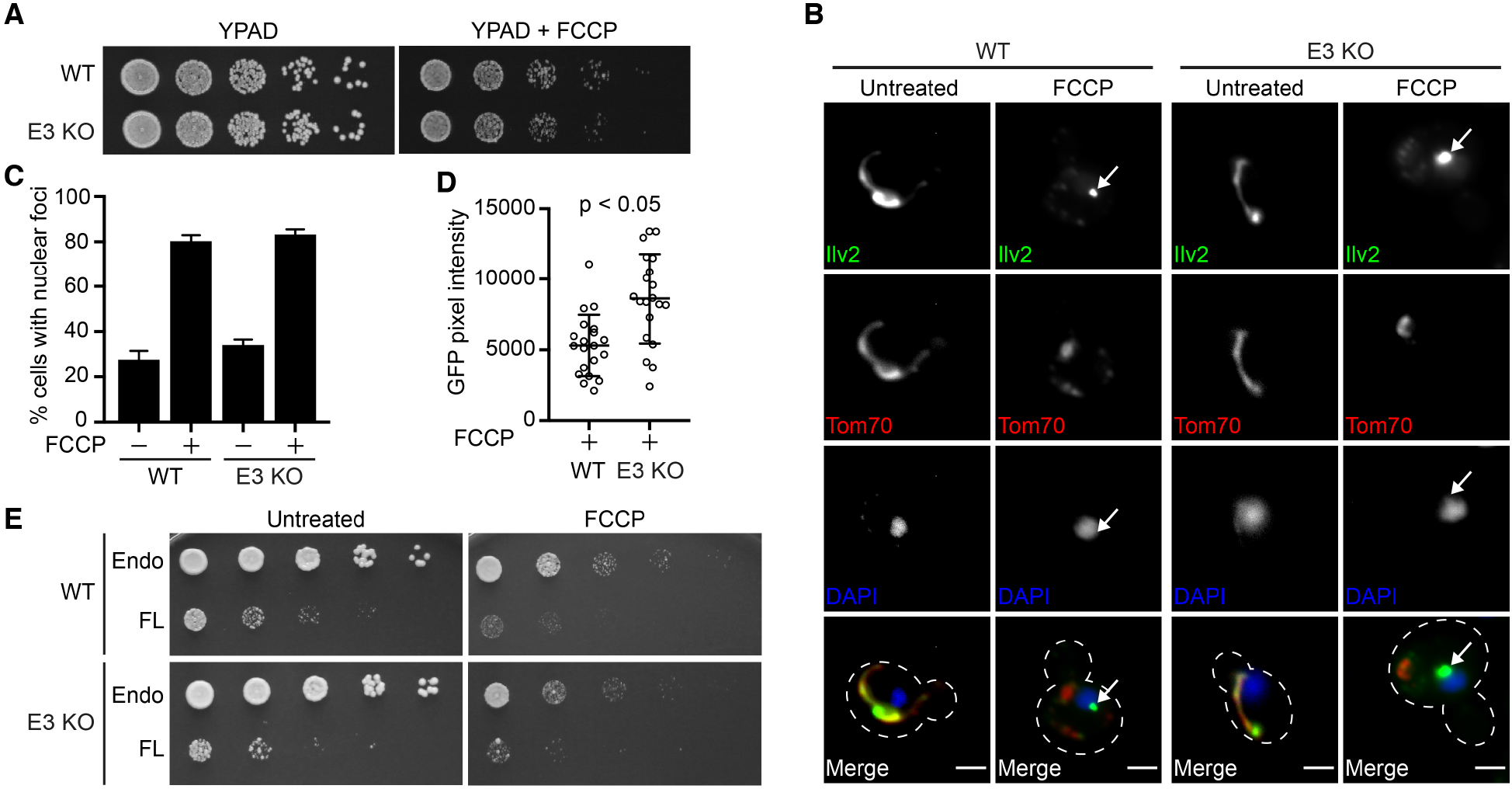
Non-imported mitochondrial precursors localize to nuclear-associated foci when clearance is impaired. (**A**) Five-fold serial dilutions of WT and E3 KO strains on YPAD −/+ FCCP agar plates. (**B**) WT and E3 KO yeast expressing Ilv2-GFP and Tom70-mCherry −/+ FCCP. Nucleus stained with DAPI. Arrows = nuclear associated foci. Bar = 2μm. (**C**) Quantification of (**B**). N > 99 cells per replicate, error bars = SEM of three replicates. (**D**) Quantification of average pixel intensity of Ilv2-GFP nuclear foci from (**B**). N=20 cells, error bars = SD, p-value = 0.0005. (**E**) Five-fold serial dilutions of WT and E3 KO strains expressing endogenous Ilv2-GFP (endo) −/+ mild overexpression of full length Ilv2-GFP (FL) from pRS413-Ilv2-GFP on SD-His −/+ FCCP agar plates.

**Fig. 4.**
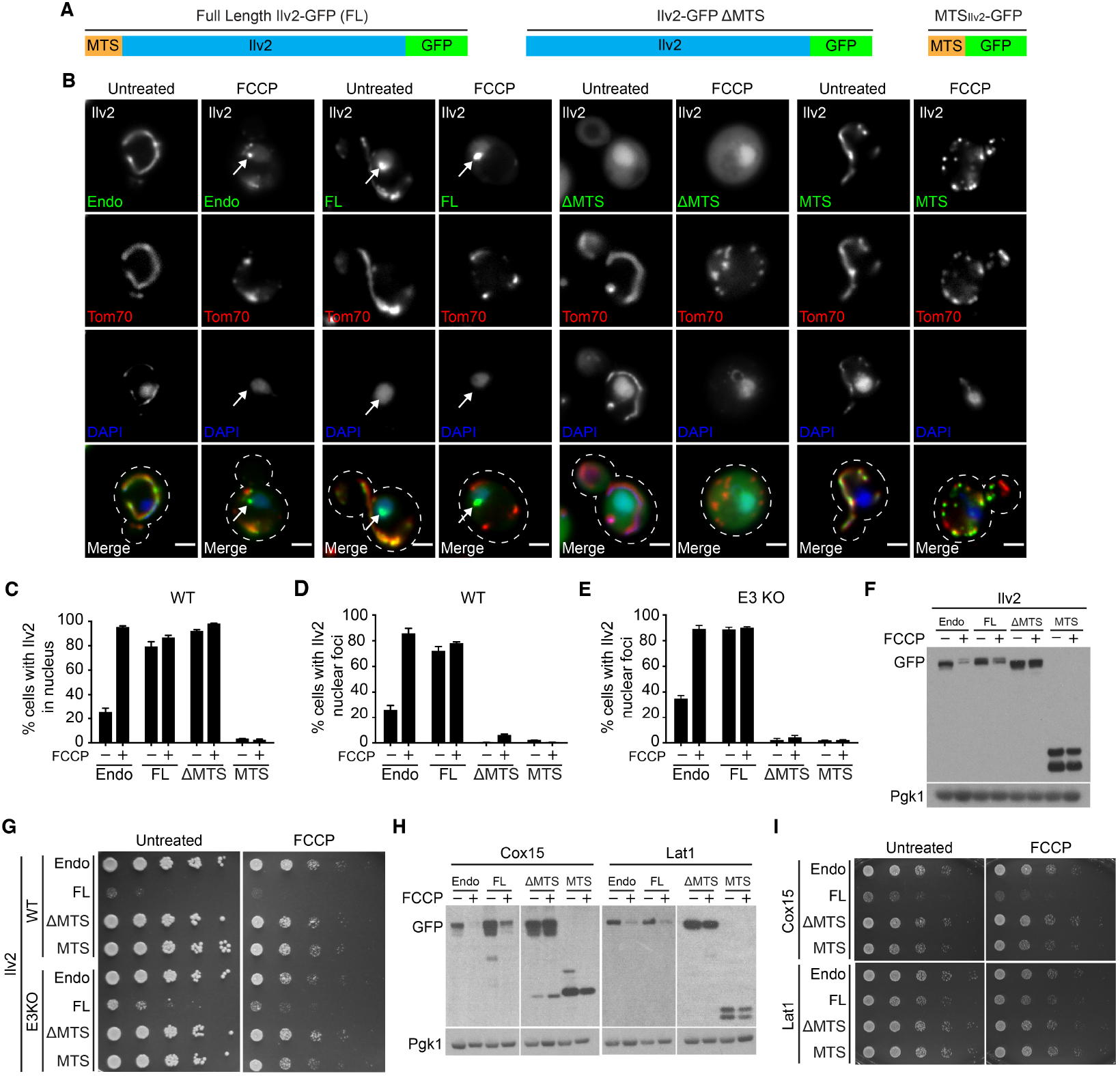
The mitochondrial targeting sequence (MTS) is required for non-imported precursor toxicity and quality control. (**A**) Schematic of full length GFP-tagged Ilv2 (FL), mitochondrial targeting sequence deleted (ΔMTS) GFP-tagged Ilv2, and MTS_Ilv2_ GFP only (MTS). (**B**) Tom70-mCherry yeast expressing endogenous Ilv2-GFP −/+ the indicated Ilv2 variant −/+ FCCP. Nucleus stained with DAPI. Arrows = nuclear associated foci. Bars = 2μm. (**C** and **D**) Quantification of cells with diffuse Ilv2 nuclear localization (C) or Ilv2 nuclear foci (D) from B. (**E**) Quantification of cells with Ilv2 nuclear foci in E3 KO strain (*san1Δ ubr1Δ doa10Δ*) conducted in parallel with (B-D). For C-E, N > 99 cells per replicate, error bars = SEM of three replicates. (**F**) Western blot of strains expressing indicated Ilv2-GFP variants −/+ FCCP. Pgk1 = loading control. (**G**) Five-fold serial dilutions of WT and E3 KO strains expressing endogenous Ilv2-GFP (endo) −/+ mild overexpression of the indicated Ilv2-GFP variants on SD-His −/+ FCCP agar plates. (**H**) Western blot on strains expressing endogenous Cox15-GFP (endo) or Lat1-GFP (endo), respectively, −/+ the mild overexpression of the indicated variants. Pgk1 = loading control. (**I**) Five-fold serial dilutions of WT strains expressing endogenous Cox15-GFP or Lat1-GFP (endo) −/+ mild overexpression of the indicated variants on SD-His −/+ FCCP agar plates.

Finally, we sought to elucidate the features of non-imported mitochondrial proteins that drive nuclear-associated aggregation, degradation and toxicity. Mitochondrial matrix proteins such as Ilv2 are synthesized as precursors with an N-terminal MTS (*19*). The MTS is removed by mitochondrial-localized proteases after import (*20*), and failure to remove and clear MTSs leads to toxicity (*31, 32*). To test whether the presence of an MTS on an unimported mitochondrial precursor protein is problematic, we analyzed strains containing single-copy plasmids expressing full-length Ilv2-GFP (FL), MTS-deleted Ilv2-GFP (ΔMTS) and MTS_Ilv2_-GFP only (MTS) from the constitutive GPD promoter (Fig. 4A). Like endogenous Ilv2-GFP (Endo), plasmid-derived FL-Ilv2-GFP localized to the nucleus and nuclear-associated foci in both wild type and E3 KO, and its abundance declined with FCCP (Fig. 4, B to F). By contrast, Ilv2 lacking an MTS (ΔMTS-Ilv2-GFP) constitutively localized to the nucleus even in the absence of FCCP, but never formed nuclear-associated foci or decreased in abundance with FCCP (Fig. 4, B to F). MTS_Ilv2_-GFP localized to mitochondria and exhibited no nuclear localization, puncta formation, or changes in total abundance with FCCP (Fig. 4, B to F). Thus, information in the mature, C-terminal portion of Ilv2 is necessary and sufficient for nuclear localization, but the presence of an MTS is required for non-imported Ilv2 degradation and sequestration into nuclear-associated foci.

Because Ilv2 lacking an MTS was not subjected to quality control, we wondered whether ΔMTS-Ilv2-GFP was still toxic to cells. In contrast to overexpressed FL Ilv2-GFP, which impaired growth of both wild type and E3 KO cells in the presence or absence of FCCP, overexpressed Ilv2 lacking its MTS did not slow cell growth, and neither did overexpressed MTS_Ilv2_-GFP alone (Fig. 4G). Thus, the presence of an MTS on unimported Ilv2 rendered the protein toxic and promoted its subsequent quality control. Importantly, the association between the presence of an MTS, degradation, and toxicity was conserved for other nuclear class proteins. Like Ilv2, Dld2 lacking its MTS constitutively localized to the nucleus, but was not subjected to degradation or sequestered into nuclear foci, and was no longer toxic to cells (fig. S4, A to F). Moreover, degradation and toxicity of Cox15 and Lat1, which are degraded by a non-nuclear proteasome pathway, also required an N-terminal MTS (Fig. 4, H and I). Thus, the presence of an MTS on an unimported mitochondrial protein drives proteotoxic stress and targets the protein for quality control.

Prior studies demonstrated that the accumulation of unimported mitochondrial precursors causes proteotoxicity (*5, 6*). To combat this stress, cells mount a coordinated response that involves upregulation of proteasome capacity (*6*, *10*), downregulation of translation (*5*) and clearance of precursors that accumulate at the mitochondrial surface (*7, 8*) and ER membrane (*9*). Here, we surveyed the mitochondrial proteome to get a clearer picture of the full spectrum of fates for unimported mitochondrial proteins. We found that mitochondrial precursors accumulate in many regions of the cell, and identified the nucleus as an important quality control destination for sequestering and destroying unimported mitochondrial proteins. Moreover, we demonstrated that the N-terminal MTS is a major driver of unimported protein toxicity. Our findings indicate that unimported mitochondrial proteins represent a large class of endogenous substrates for nuclear protein quality control. This discovery raises the intriguing possibility that unimported mitochondrial proteins may synergize with other aggregate-prone proteins to overwhelm protein quality control systems during aging and disease (*33*). Future studies to determine what drives unimported mitochondrial proteins to various cellular destinations, and elucidate the coordination between unimported mitochondrial quality control pathways will help illuminate how cells cope with the proteotoxic burden that arises during times of mitochondrial dysfunction.

## Acknowledgements

We thank members of the A.L.H. laboratory and Janet Shaw (Utah) for discussion and manuscript comments, Tom Tedeschi (Utah) for technical assistance, Dr. Nikolaus Pfannner for Tim44 and Tom70 antisera and Dr. Toshiya Endo for Tom40 antisera.

## Funding

Research was supported by NIH grants AG043095 and GM119694, (A.L.H.). A.L.H. was further supported by an American Federation for Aging Research Junior Research Grant, United Mitochondrial Disease Foundation Early Career Research Grant, Searle Scholars Award, and Glenn Foundation for Medical Research Award.

## Author Contributions

All authors conceived aspects of the project, designed experiments, and discussed and analyzed results. V.P.S.S., W.A.B., T. X., and C.S.K. conducted experiments. V.P.S.S. and A.L.H. wrote and edited the manuscript.

## Competing interests

Authors declare no competing interests.

## Data and materials availability

All data is available in the main text or the supplementary materials

## Supplementary Materials

### Materials and Methods

#### Reagents

Chemicals were obtained from the following sources: β-Estradiol (E8875), Carbonyl cyanide 4-(trifluoromethoxy) phenylhydrazone (C2920), cOmplete Protease Inhibitor Cocktail (11697498001), dimethyl sulfoxide (D2650), Cycloheximide (C1988), Doxycycline hyclate (C9891), polyvinylpyrrolidone (PVP40), Pepstatin (10253286001), Phenylmethylsulfonyl fluoride (P7626), calcofluor Fluorescent Brightener 28 (F3543) from Millipore Sigma, 4’,6-Diamidino-2-Phenylindole Dihydrochloride (DAPI) (D130), ProLong™ Glass Antifade Mountant with NucBlue™ Stain (P36981) from ThermoFisher, (S)-MG-132 (10012628) from Cayman Chemical, N-Ethylmaleimide (NEM) (S3876), IGEPAL NP-40 (CA-630) from Sigma-Aldrich, Zymolyase 100T (Z1004) from US Biological Life Sciences, Triton X-100 (1610407) from Biorad, Paraformaldehyde (100503-914) from VWR, and Dithiothreitol (DTT10) from GOLDBIO. Antibodies and other reagents are described in the appropriate section below.

#### Yeast Strains

All yeast strains are derivatives of Saccharomyces cerevisiae S288c (BY) (34) and are listed in Supplementary Table 2. Strains expressing fluorescently tagged proteins from their native loci were created by one step PCR-mediated C-terminal endogenous epitope tagging using standard techniques and the oligo pairs listed in Supplementary Table 3 (34, 35). Plasmid template for GFP and mCherry tagging was from the pKT series of vectors (35), plasmid template for RITE tagging was previously described pVL015 (21), and plasmid templates for FLAG, HA, and mCherry tagging were pFA6A-5FLAG-KanMX (Addgene 15983) (36), pFA6A-3HA-His3MX (Addgene 41600) (37), pFA6A-3HA-KanMX (Addgene 39295) (38), and pFA6A-mCherry-HphMX (Addgene 105156) (39). Deletion strains were created by one step PCR-mediated gene replacement using the oligos pairs listed in Supplementary Table 3 and plasmid templates of pRS series vectors (34). Correct integrations were confirmed with a combination of colony PCR across the chromosomal insertion site and correctly localized expression of the fluorophore by microscopy. The strain collection used for screening in Figure 1 expressed Tom70-mCherry/any protein-GFP and was created previously (23). The genotype of all strains in the collection is MATa/MATαhis3Δ1/his3Δ1 leu2Δ0/leu2Δ0 ura3Δ0/ura3Δ0 met15Δ0/+ lys2Δ0/+ anygene-GFP-His3MX/+ TOM70-mCherry-KanMX/+.

#### Yeast Cell Culture and Media

For all microscopy and western blot experiments, yeast were grown exponentially for 15 hours up to a maximum density of 1 × 10^7^ cells/ml prior to starting any treatments. Cells were cultured as indicated in the Main Text and Figure Legends in YPAD medium (1% yeast extract, 2% peptone, 0.005% adenine, 2% glucose) or synthetic defined medium lacking histidine (SD-His) (0.67% yeast nitrogen base without amino acids, 2% glucose, supplemented nutrients 0.074 g/L each adenine, alanine, arginine, asparagine, aspartic acid, cysteine, glutamic acid, glutamine, glycine, myo-inositol, isoleucine, lysine, methionine, phenylalanine, proline, serine, threonine, tryptophan, tyrosine, uracil, valine, 0.369 g/L leucine, 0.007 g/L para-aminobenzoic acid). FCCP and MG-132 were used at a final concentration of 10 μM and 50 nM respectively. All FCCP and/or MG-132 treatments were conducted for six hours. For knockdown of TOM40 expressed under control of the tetracycline promoter, cultures were grown in log-phase for 16 hours in the presence of doxycycline (20 μg/mL) prior to any experimental treatments. The wild-type control strain was cultured under the same conditions. For RITE tag-switching experiments, β-Estradiol was added to cultures at a final concentration of 1 μM to induce tag switching. FCCP was added to cultures at a final concentration of 10 μM at the same time of β-Estradiol. Cultures were imaged after 6 hours of treatment.

#### Plasmids and Cloning

Centromeric yeast plasmids expressing GPD-promoter driven full-length, MTS-deleted, or MTS-only versions of Ilv2, Lat1, Cox15, and Dld2 fused to C-terminal GFP epitopes were assembled using Gibson Assembly^®^Master Mix (E2611L, NEB) following the manufacturer’s instructions. Plasmid names and construction details (including PCR templates, oligo pairs, and digested plasmid templates) used in Gibson Assembly are listed in Supplementary Table 4. PCR amplifications from yeast genomic DNA and plasmid DNA were conducted with Phusion Polymerase (M0530L, NEB) using oligonucleotides listed in Supplementary Table 3. Plasmids were verified by sequencing.

#### MTS Prediction

Mitochondrial targeting sequences for Ilv2, Cox15, Lat1, and Dld2 were predicted using Mitoprot (40). Correct MTS prediction was confirmed by analyzing localization of C-terminal GFP-tagged versions of MTS-only or MTS-deleted proteins via microscopy.

#### Microscopy

200-300 nm optical Z-sections of live yeast cells were acquired with an AxioImager M2 (Carl Zeiss) equipped with an Axiocam 506 monochromatic camera (Carl Zeiss) and 100× oil-immersion objective (Carl Zeiss, Plan Apochromat, NA 1.4), or with an AxioObserver 7 (Carl Zeiss) equipped with a PCO Edge 4.2LT Monochrome, Air Cooled, USB 3 CCD camera with a Solid-State Colibri 7 LED illuminator and 100X oil-immersion objective (Carl Zeiss, Plan Apochromat, NA 1.4). All images were acquired with ZEN (Carl Zeiss), and processed with Fiji (NIH). All images shown in Figures represent a single optical section.

#### DAPI staining

Yeast cells were stained with DAPI by incubating cultures for ten minutes in respective growth media with DAPI (1μg/ml).

#### Quantification of Nuclear-Associated Foci Intensity

Mean GFP pixel intensity of nucleus and nuclear associated foci was calculated via line scan analysis of pixel intensity from maximum intensity projections on 20 cells using FIJI (NIH) (41). Nucleus stained by DAPI was used as a reference to draw lines of ~2.5 μm for analysis.

#### Determination of Replicative Age

Yeast strains exponentially growing for 15 hours up to a maximum density of 1 × 10^7^ cells/ml were stained with for 5 minutes in YPAD with 5 μg/ml of Fluorescent Brightener 28 (F3543, Millipore Sigma), which stains bud scars. The replicative age of each yeast cell was determined by counting of the number of bud scars after staining. Cells with less than five bud scars were categorized as young and cells with five or more bud scars were categorized as old.

#### Indirect Immunofluorescence (IIF) Staining

For IIF staining, cells were harvested by centrifugation and fixed in 10 ml fixation medium (4% Paraformaldehyde in YPAD) for 1 hour. Fixed yeast cells were washed with Wash Buffer (0.1M Tris, pH=8, 1.2M Sorbitol) twice and incubated with DTT (10mM DTT in 0.1M Tris, pH=9.4) for 10 min. Spheroplasts were generated by incubating cells in solution containing 0.1M KPi, pH=6.5, 1.2M Sorbitol and 0.25mg/ml Zymolyase at 30°C for 30 minutes. Spheroplasts were gently diluted in 1:40 using Wash Buffer and attached to glass slides pre-coated with 0.1% poly-L-Lysine (2mg/ml). Samples were permeabilized in cold 0.1% Triton-X100 in PBS for 10 min at 4°C, briefly dried and blocked (30 min at room temperature) in Wash Buffer containing 1% BSA. After blocking, samples were incubated with 1:200 diluted anti-FLAG primary antibody (F1804, Millipore Sigma) for 90 minutes followed by washing 10 times. Samples were then incubated with 1:300 diluted secondary antibody (A32723, Invitrogen) followed by washing 10 times. Antibody dilutions were made using Wash Buffer containing 1% BSA. Samples were washed with Wash Buffer containing 1% BSA and 0.1% Tween-20. Slides were washed twice with Wash Buffer before sealing, and mounted with hardset medium containing NucBlue™ stain (P36981, Invitrogen) overnight. Widefield images were acquired as described above in microscopy section.

#### Protein Preparation and Western Blotting

Western blotting of yeast extracts was carried out as described previously(23). Briefly, 1 × 10^7^ log phase yeast cells were harvested and resuspended in 50 μl of H_2_O. 50 μl of NaOH (1 M) was added to cell suspension and incubated for 5 minutes at room temperature. Cells were centrifuged at 20,000xg for 10 min at 4°C and cell pellets were resuspended in SDS lysis buffer (30 mM Tris-HCl pH 6.8, 3% SDS, 5% glycerol, 0.004% bromophenol blue, 2.5% β-mercaptoethanol). Cells extracts were resolved on Bolt 4-12% Bis-Tris Plus Gels (NW04125BOX, Thermo Fisher) with NuPAGE MES SDS Running Buffer (NP0002-02, Thermo Fisher) and transferred to nitrocellulose membranes. Membranes were blocked and probed in blocking buffer (1XPBS, 0.05% Tween 20, 5% non-fat dry milk) using the primary antibodies for GFP (1814460001, Sigma Millipore) or Pgk1 (22C5D8, abcam), and HRP conjugated secondary antibodies (715-035-150, Jackson Immunoresearch). Blots were developed with SuperSignal West Pico Chemiluminescent substrate (34580, Thermo Fisher) and exposed to films. Blots were developed using film processor (SRX101, Konica Minolta) or a Chemidoc MP system (BioRad).

#### Nuclear Enrichment

Cells were grown in log-phase overnight as described above followed by treatment with MG-132 and −/+ FCCP for 4 hours. 4 × 10^8^ total cells were harvested. Cells were washed with ddH_2_O, and the wet weight of the pellet was recorded. Cells were incubated in DTT Buffer (100 mM Tris-HCl pH 9.5, 10 mM DTT) and 50 nM MG-132 with gentle shaking at 30°C for 20 min. Cells were then spheroplasted via incubation in Zymolyase Buffer (1.2 M sorbitol, 20 mM K_2_HPO_4_, pH 7.4), 50 nM MG-132, and 1 mg of Zymolyase 100T (Z1004, US Biological Life Sciences) per 1 g cell pellet for 1 hour at 30°C with gentle shaking. Spheroplasts were washed once with Zymolyase Buffer, and then all subsequent steps were carried out on ice. Spheroplasts were dounce-homogenized with 35 strokes in 5 mL of polyvinylpyrrolidone-40 solution (8% PVP-40, 20 mM K-phosphate, 7.5 μM MgCl_2_, pH 6.5), 0.025% Triton X-100, 5 mM DTT, 50 μL Solution P (20 mg/mL PMSF, 0.4 mg/mL Pepstatin A in ethanol), and 50 μL 100X cOmplete protease inhibitor cocktail (11697498001, Millipore Sigma). Next, 15 mL of PVP-40 solution, 15 μL Solution P, and 15 μL PIC was added, and spheroplasts were dounce-homogenized with an additional 5 strokes. PVP-40 ensures nuclei stay intact during lysis (42). The cell lysate was centrifuged for 3000 × g for 5 min. The resulting supernatant was discarded, and pellets were washed once and resuspended in 1ml of IP Buffer (50mM Tris pH7.5, 150mM NaCl, 1mM EDTA, 10% Glycerol, 1% IGEPAL (NP-40 substitute), 100uM PMSF). Intact nuclei, which are more resistant to NP-40 than other cellular membranes, were immobilized non-specifically to magnetic agarose beads (BMAB 20, Chromotek) via incubation at 4°C for 2-3 hr. After binding, nuclei were washed 4 × 15 min in IP buffer at 4°C. Nuclear-enriched extracts were eluted by incubating beads in 2X Laemmli buffer (63 mM Tris pH 6.8, 2% (w/v) SDS, 10% (v/v) glycerol, 1 mg/ml bromophenol blue, 1% (v/v) b-mercaptoethanol) at 90°C for 10 minutes. Eluates were subjected to SDS-PAGE and Western Blotting with primary anti-HA antibody (11583816001, Sigma Millipore), anti-Tom70 and Tim44 antisera (gifts from Dr. Nikolaus Pfanner, University of Freiburg), anti-GFP antibody (1814460001, Millipore Sigma) and anti-H2b antibody (39947, Active Motif). Effectiveness of nuclear enrichment was indicated by increase in relative abundance of nuclear markers H2B and Nup49-GFP, and decrease in Tom70 and Tim44 in nuclear extracts compared to whole cell lysate. Nuclei were monitored during isolation by visualizing Nup49-GFP via fluorescence microscopy.

#### Cycloheximide-Chase Analysis

Exponentially growing cells were treated −/+ FCCP for 4 hours, after which, cycloheximide (100 μg/ml) was added to the cultures. The time zero sample was collected immediately after adding cycloheximide. For all other time-points, samples were collected by harvesting an equal volume of media to that which was harvested at time zero. Samples were then subjected to SDS-PAGE and Western Blotting with primary antibodies for HA (11583816001, Sigma Millipore) or GFP (1814460001, Sigma Millipore) and Pgk1 (22C5D8, abcam). Blots were developed as described above.

#### Microscopy and Western Blot Screens

Individual strains listed in Supplementary Table 1 from the Tom70-mCherry/mitochondrial protein GFP collection were cultured in batches overnight in YPAD as described above and then incubated +/− FCCP for six hours. After treatment, cultures were split for simultaneous microscopy and western Blot analysis. Images and western blots were analyzed and scored by three independent researchers. A subset of strains from each class was reconstructed and reanalyzed with both FCCP and genetic ablation of mitochondrial import. Class assignments were based on combined results of microscopy and western blot analysis and were as follows: Class 1 (nucleus), small to large decrease in protein levels and localized to the nucleus in the presence of FCCP; Class 2 (mitochondria), minimal change in protein level and robustly localized to mitochondria in the presence of FCCP; Class 3 (cytoplasm), no change or an increase in protein level and localized predominantly to the cytoplasm with FCCP treatment; Class 4 (ER), mild or no change in protein abundance and localized to ER upon FCCP; Class 5 (reduced abundance), large reduction in protein abundance and no longer easily detectable via microscopy with FCCP treatment.

#### Immunoprecipitation

Cells were grown as described above and treated +/− FCCP and MG-132 for six hours. 1 × 10^8^ total cells were harvested, resuspended in 1ml of lysis Buffer (50mM Tris pH7.5, 150mM NaCl, 1mM EDTA, 10% Glycerol, 1% IGEPAL (NP-40 substitute), 100uM PMSF and 10mM NEM and lysed with glass beads using an Omni Bead Ruptor 12 Homogenizer (8 cycles of 20 seconds each). Cells lysates were cleared by centrifugation at 20000g and supernatant was moved to a new tube. Cell pellets were resuspended in 50 μl of SUME buffer (1% SDS, 8 M Urea, 10 mM MOPS, pH 6.8, 10 mM EDTA and 10mM NEM) and heated at 55 °C for 5 minutes. 50 μl of cell pellet resuspension was combined with supernatant from lysate clearance centrifugation and total volume was adjusted to 1ml by adding lysis buffer. Lysates were incubated with 25 μl of anti-GFP bead slurry (GTMA, GFP-Trap®_MA, chromotek) at 4°C for 3-4 h and then washed 4X for 10 min each in lysis buffer (without NEM). Immunoprecipitated proteins were eluted by incubating beads in 2X Laemmli buffer (63 mM Tris pH 6.8, 2% (w/v) SDS, 10% (v/v) glycerol, 1 mg/ml bromophenol blue, 1% (v/v) b-mercaptoethanol) at 90°C for 10 minutes. Eluates were subjected to SDS-PAGE and Western Blotting with primary anti-ubiquitin antibody (PA1-187, ThermoFisher) and anti-GFP antibody (1814460001, Sigma Millipore). Blots were developed as described above.

#### Statistics

Experiments were repeated at least three times and all attempts at replication were successful. For all quantifications, number of cells scored is included in Figure Legends. Differences in means were compared using two-tailed t-tests at the 5% significance level. No randomization or blinding was used in experiments. All analysis was done with GraphPad Prism version 8.01.

---

**Fig. S1.**
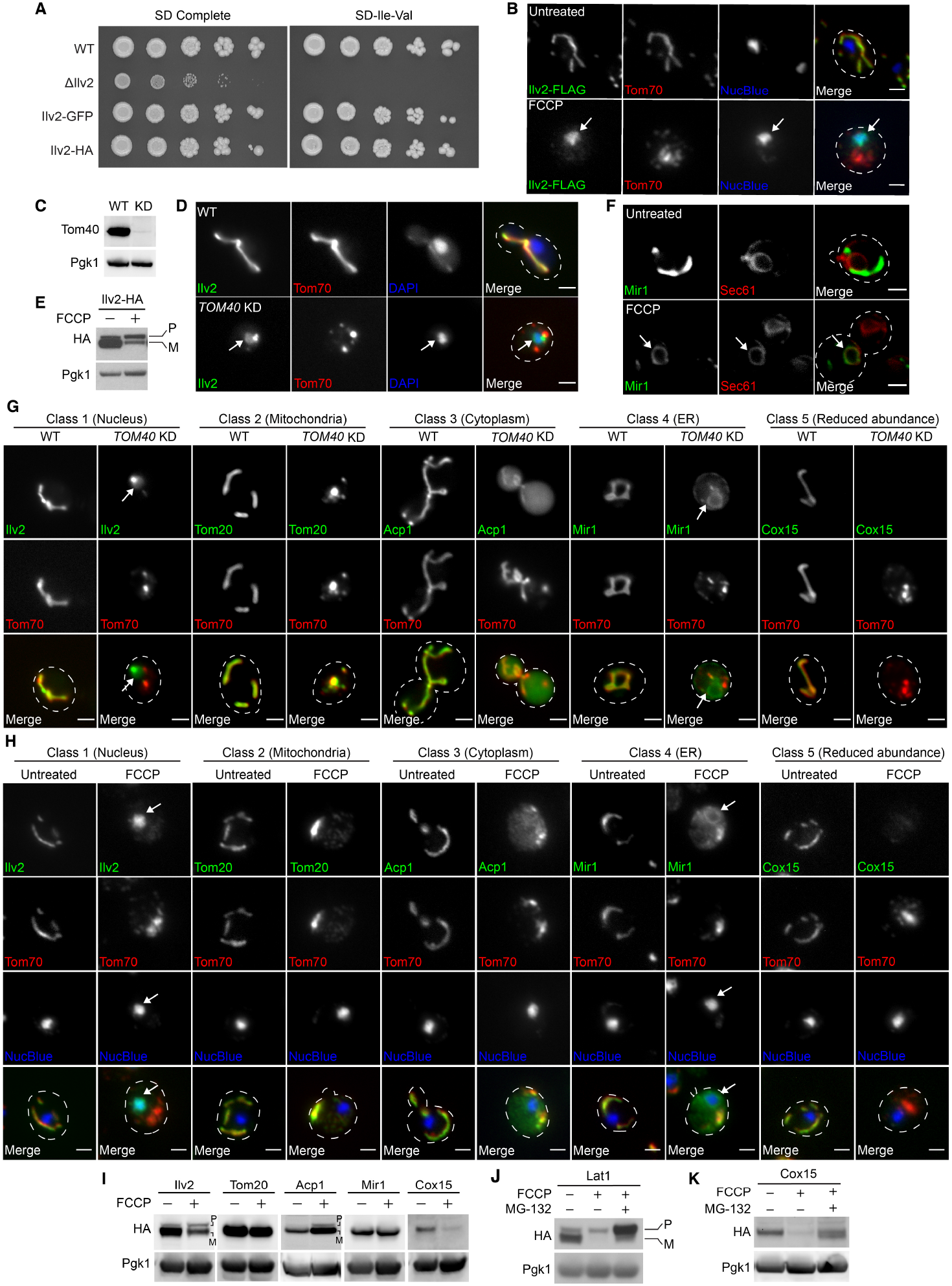
The nucleus is one of several non-imported mitochondrial precursor protein fates. (**A**) Five-fold serial dilutions of WT, Ilv2 KO, and GFP or HA tagged Ilv2 yeast strains on SD complete or isoleucine and valine dropout agar plates. (**B**) Indirect immunofluorescence of yeast expressing Ilv2-FLAG and Tom70-mCherry. (**C**) Western blot for Tom40 in wild type (WT) and *tet_p_-TOM40* (*KD*) strains in the presence of doxycycline. (**D**) Tom70-mCherry wild type (WT) and *tet_p_-TOM40* (*TOM40 KD*) yeast expressing Ilv2-GFP in the presence of doxycycline. (**E**) Western blot of yeast expressing Ilv2-HA −/+ FCCP. (**F**) Yeast expressing ER marker Sec61-mCherry and Mir1-GFP −/+ FCCP. (**G**) Tom70-mCherry wild type (WT) and *tet_p_-TOM40* (*TOM40 KD*) yeast expressing the indicated GFP-tagged mitochondrial proteins in the presence of doxycycline. (**H**) Indirect immunofluorescence of yeast expressing the indicated FLAG-tagged proteins and Tom70-mCherry. (**I** to **K)** Western blots of yeast expressing the indicated HA-tagged mitochondrial proteins −/+ FCCP (H) or −/+ FCCP −/+ MG-132 (**I-J**). P = precursor form, M = mature form. Pgk1 = loading control. In **B, D**, and **F-H**, bar = 2μm. Nucleus in (**B**, **D**, and **H)** stained with NucBlue or DAPI. Arrows indicate nucleus (**B, D**, and **G-H**, class 1) or ER (**F and G-H, and H**, class 4).

**Fig. S2.**
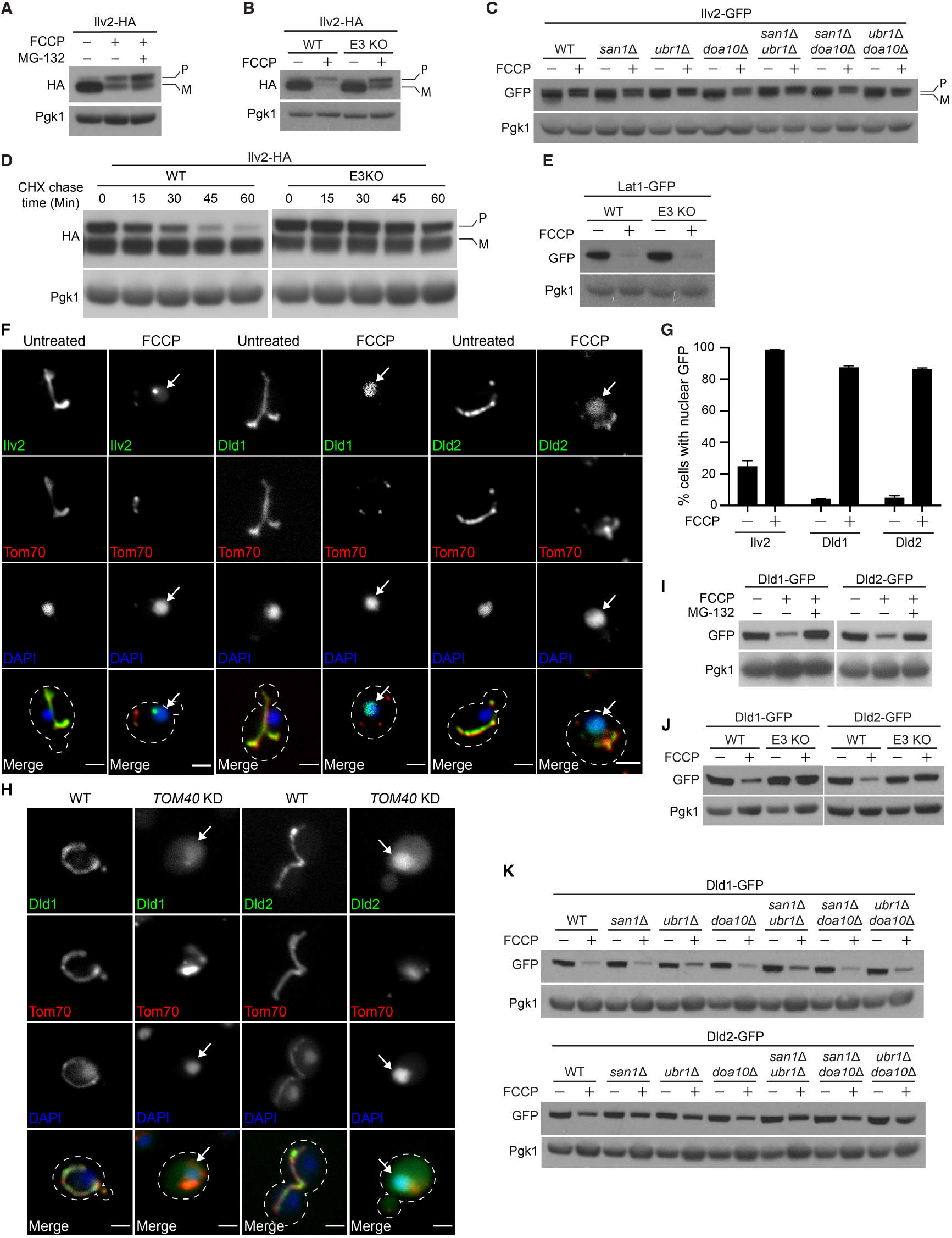
Nuclear protein quality control promotes unimported mitochondrial protein degradation. (**A**) Western blot of yeast expressing Ilv2-HA −/+ FCCP −/+ MG-132. (**B**) Western blot of yeast expressing the Ilv2-HA −/+ FCCP in WT and E3 KO strains. (**C**) Western blot of yeast expressing Ilv2-HA −/+ FCCP in WT and the indicated mutant yeast strains. (**D**) Western blots showing the CHX chase of Ilv2-HA in WT and E3 KO strains in the presence of FCCP. (**E**) Western blots of yeast expressing the Lat1-GFP −/+ FCCP in WT and E3 KO strains. (**F**) Yeast expressing the indicated GFP and mCherry tagged mitochondrial proteins −/+ FCCP. (**G**) Quantification of (**F**). N > 99 cells per replicate, error bars = SEM of three replicates. (**H**) Tom70-mCherry wild type (WT) and *tet_p_-TOM40* (*TOM40 KD*) yeast expressing the indicated GFP-tagged mitochondrial proteins in the presence of doxycycline. (F, H) Nucleus stained with DAPI, Arrows = nucleus. Bar = 2μm. (**I**)Western blots of yeast strains expressing indicated GFP-tagged mitochondrial proteins −/+ FCCP −/+ MG-132. (**J** and **K**) Western blots of yeast expressing the indicated GFP-tagged mitochondrial proteins −/+ FCCP. Pgk1 = loading control. (**B, D, E**, and **J**) E3 KO = *san1Δ ubr1Δ doa10Δ*. (**A**-**D)**, P = precursor, M = mature. Pgk1 = loading control.

**Fig. S3.**
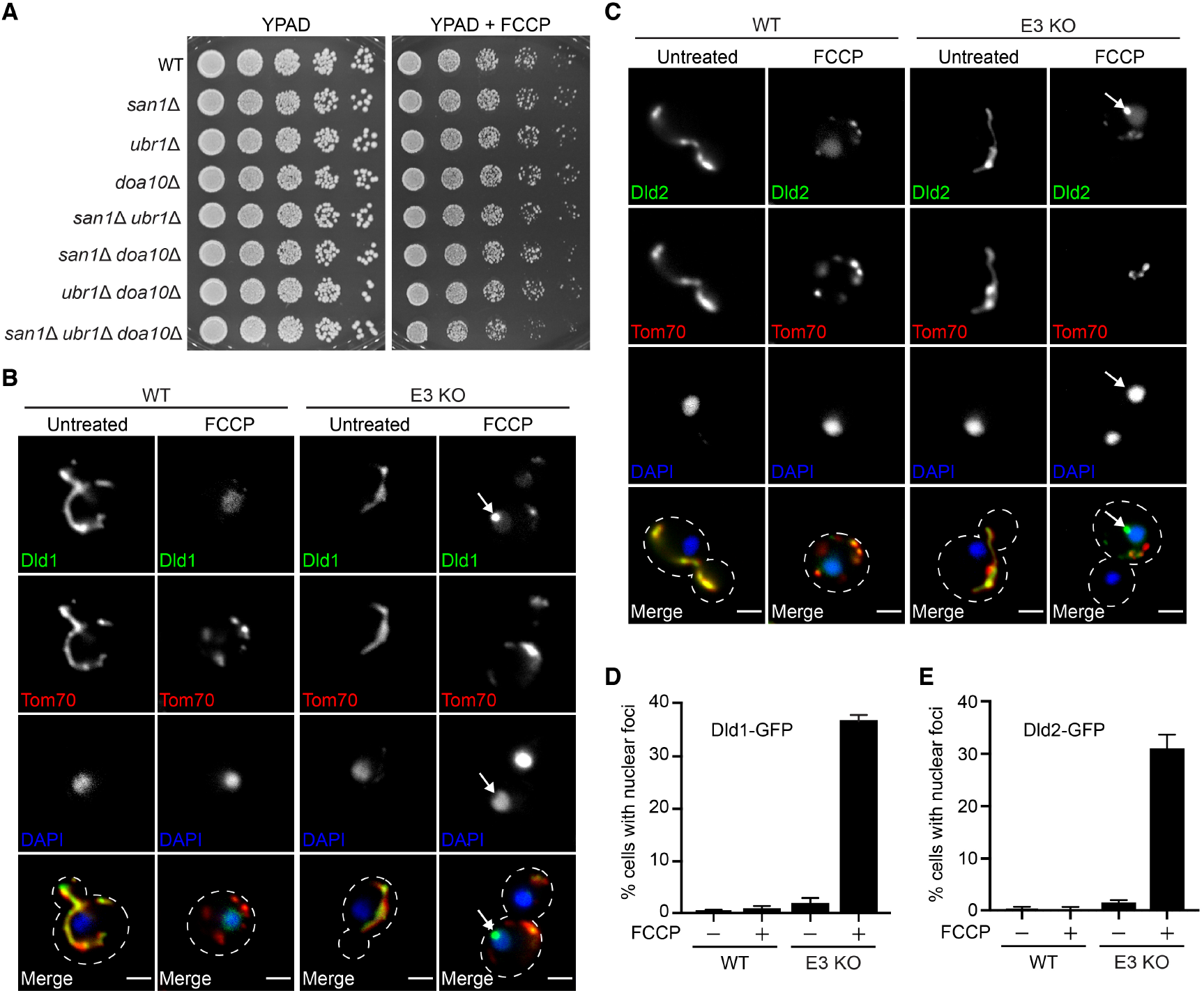
Impaired clearance of non-imported mitochondrial proteins targets them to nuclear associated foci. (**A**) Five-fold serial dilutions of WT and the indicated mutant strains on YPAD −/+ FCCP agar plates. **(B** and **C**), WT and E3 KO (*san1Δ ubr1Δ doa10Δ*) yeast expressing Dld1-GFP or Dld2-GFP and Tom70-mCherry −/+ FCCP. Nucleus stained with DAPI, arrows = nuclear associated foci, and bar = 2μm. (**D** and **E**) Quantification of (**B**) and (**C**), respectively. N > 99 cells per replicate, error bars = SEM of three replicates.

**Fig. S4.**
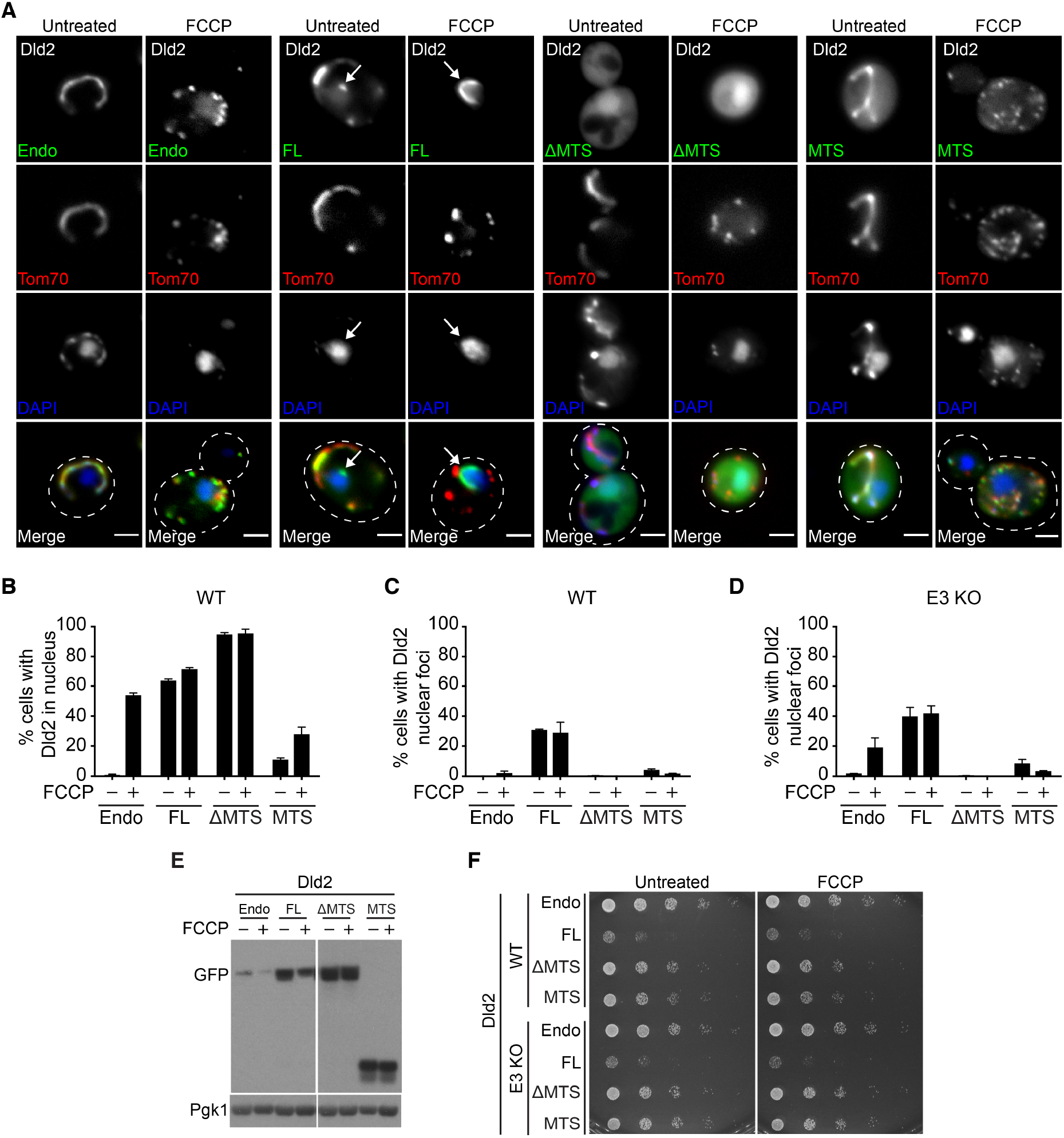
The MTS is required for non-imported precursor toxicity and degradation. (**A**) Tom70-mCherry yeast expressing endogenous Dld2-GFP −/+ the indicated Dld2 plasmid-expressed variant −/+ FCCP. Nucleus stained with DAPI. Arrows = nuclear associated foci. Bars = 2μm. (**B** and **C**) Quantification of cells with diffuse Dld2-GFP nuclear localization (**B**) or Dld2-GFP nuclear foci (**C**) from (**A**). (**D**) Quantification of cells with Dld2-GFP nuclear foci in E3 KO strain (*san1Δ ubr1Δ doa10Δ*) conducted in parallel with (**A-C)**. For (**B-D)**, N > 99 cells per replicate, error bars = SEM of three replicates. (**E**) Western blot of strains expressing indicated Dld2-GFP variants −/+ FCCP. Pgk1 = loading control. (**F**) Five-fold serial dilutions of WT and E3 KO strains expressing endogenous Dld2-GFP (endo) −/+ mild overexpression of the indicated Dld2-GFP variants on SD-His −/+ FCCP agar plates.

**Table S1.**
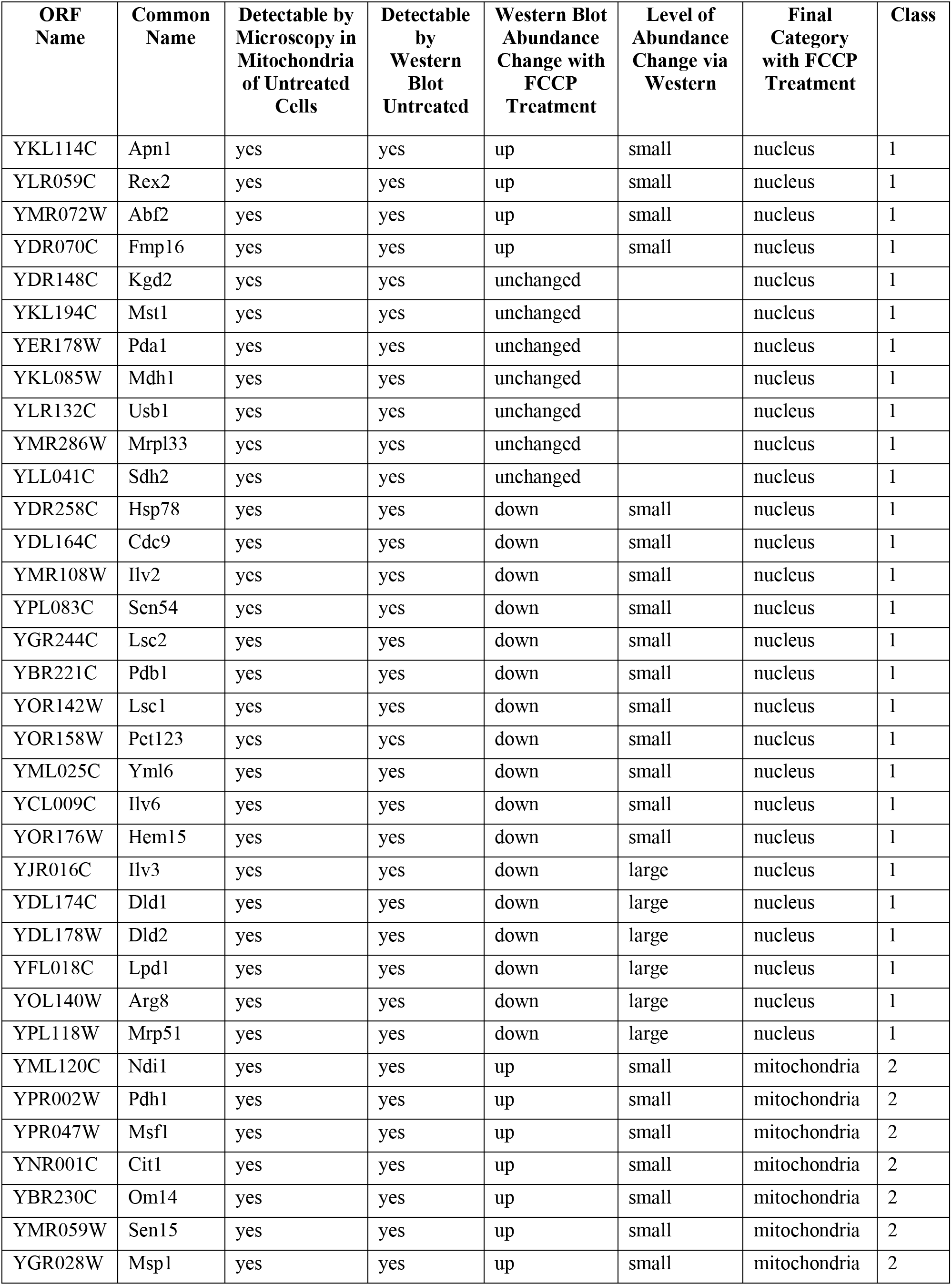
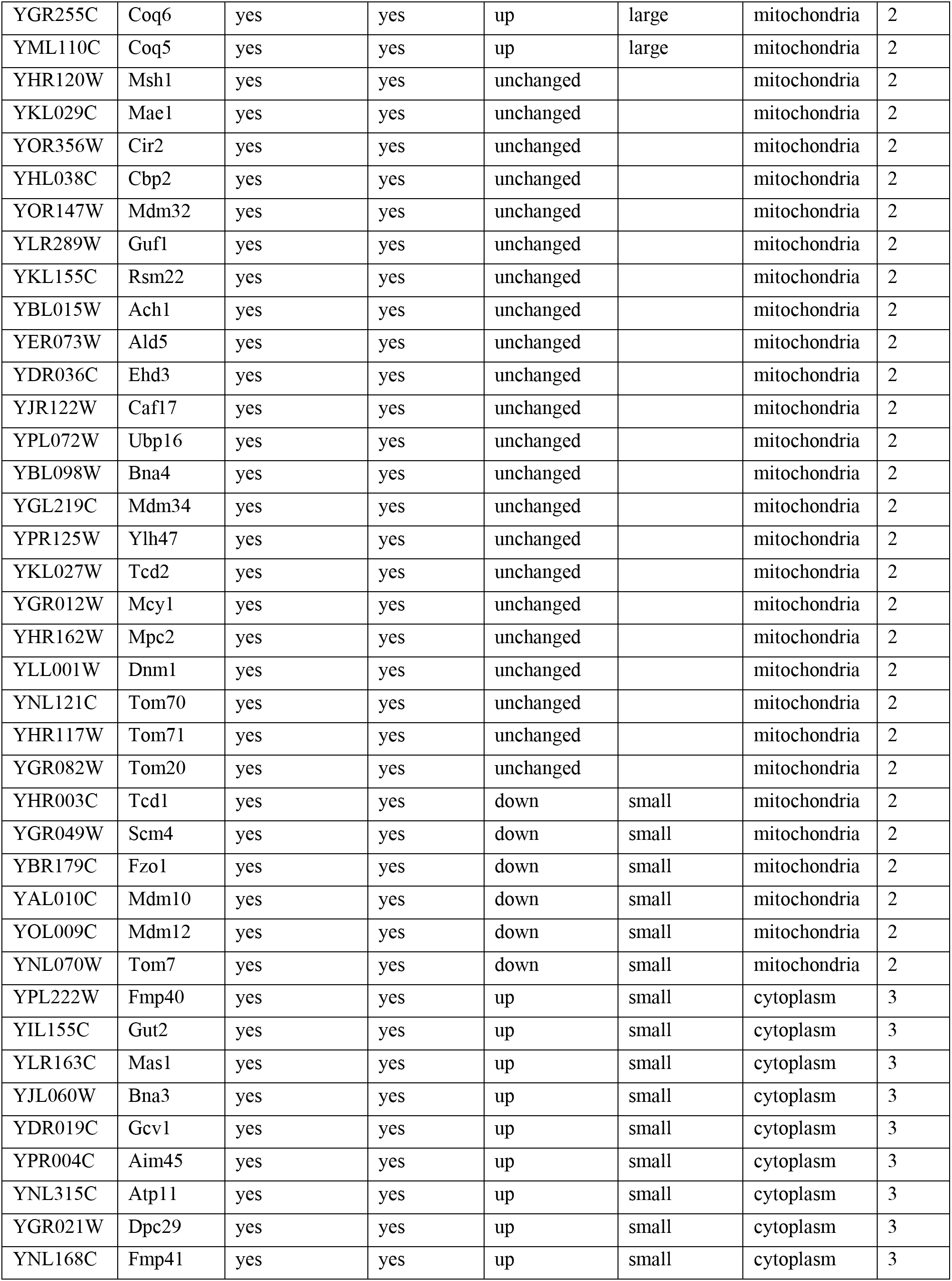
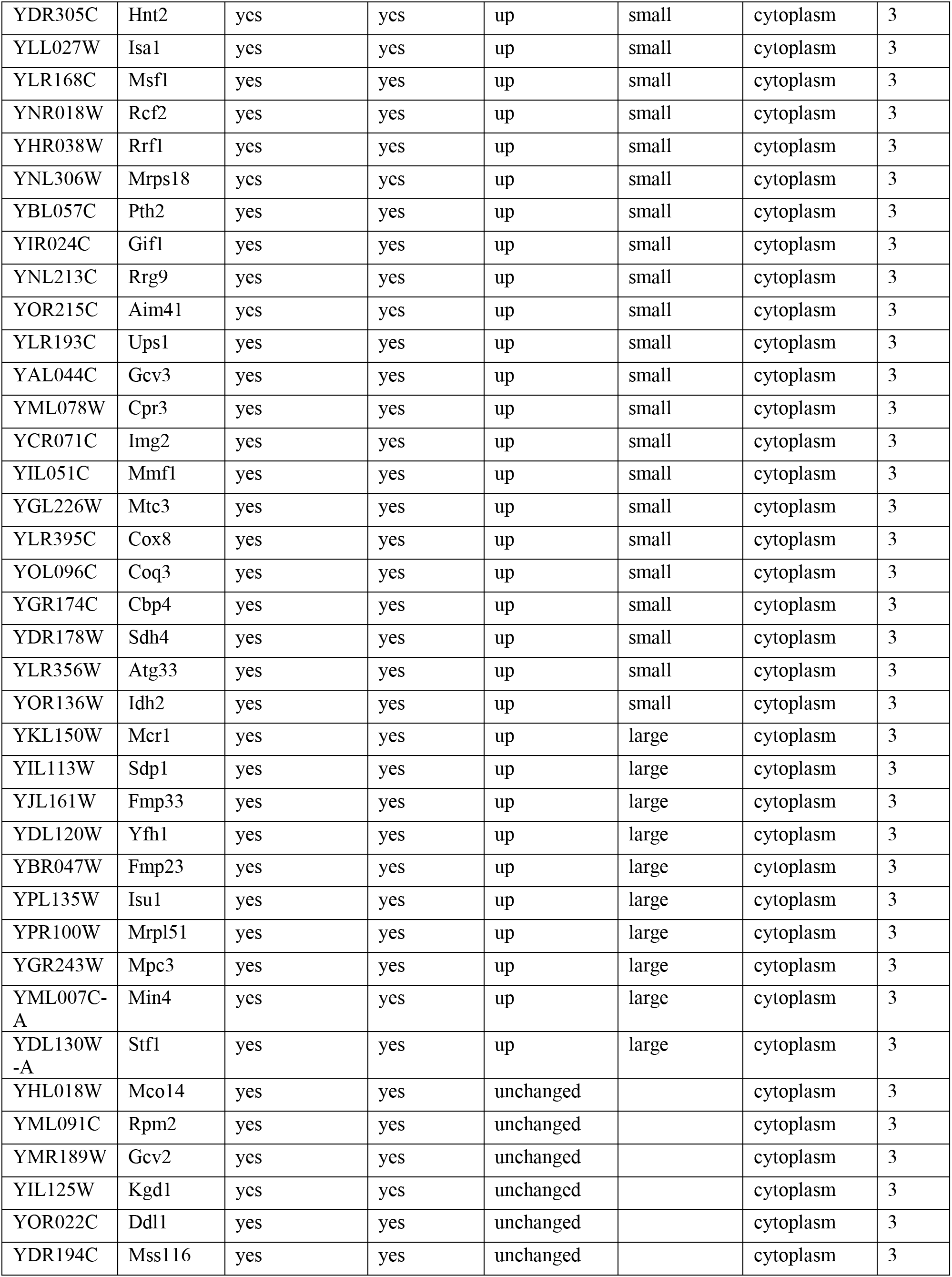
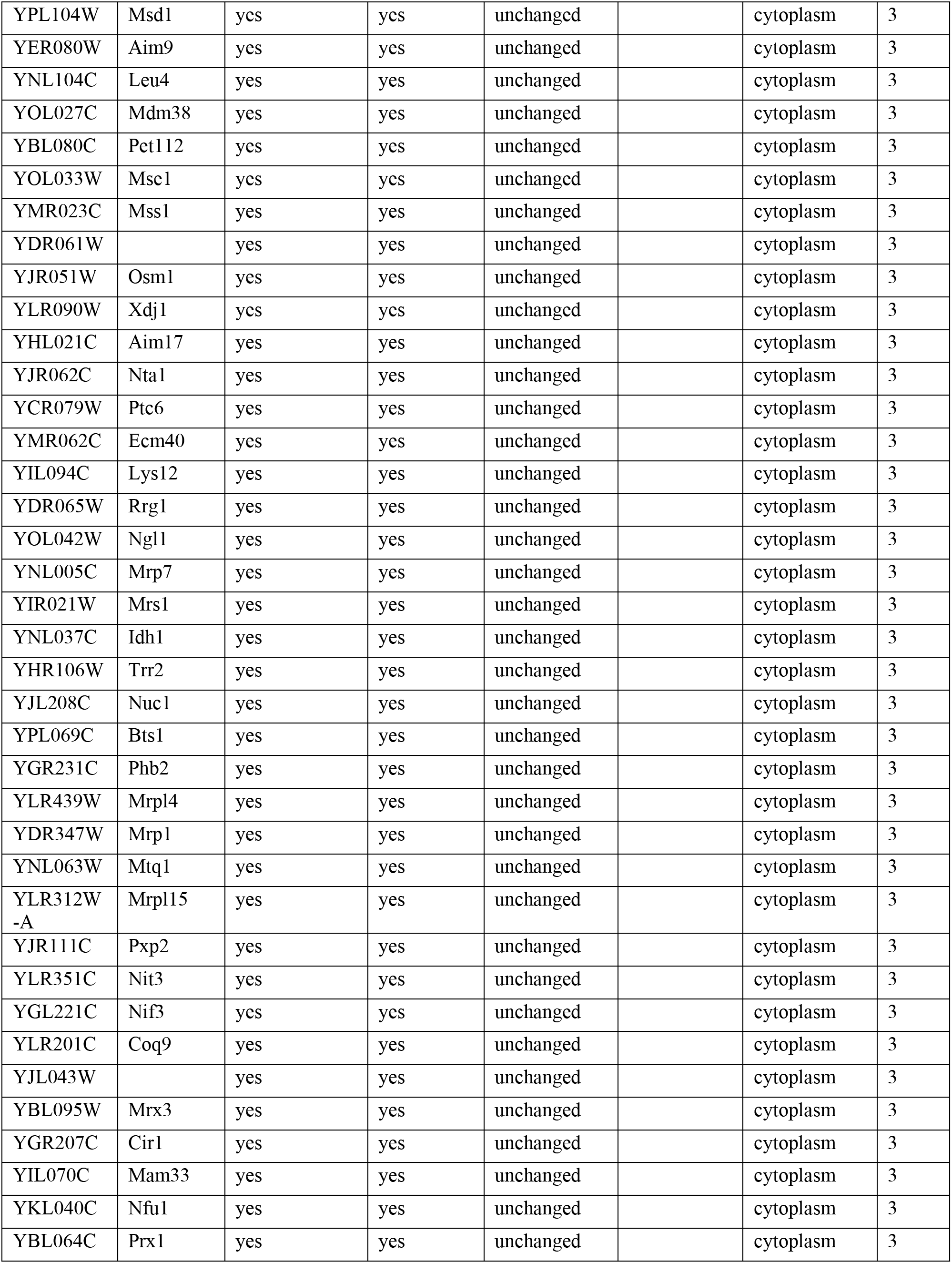
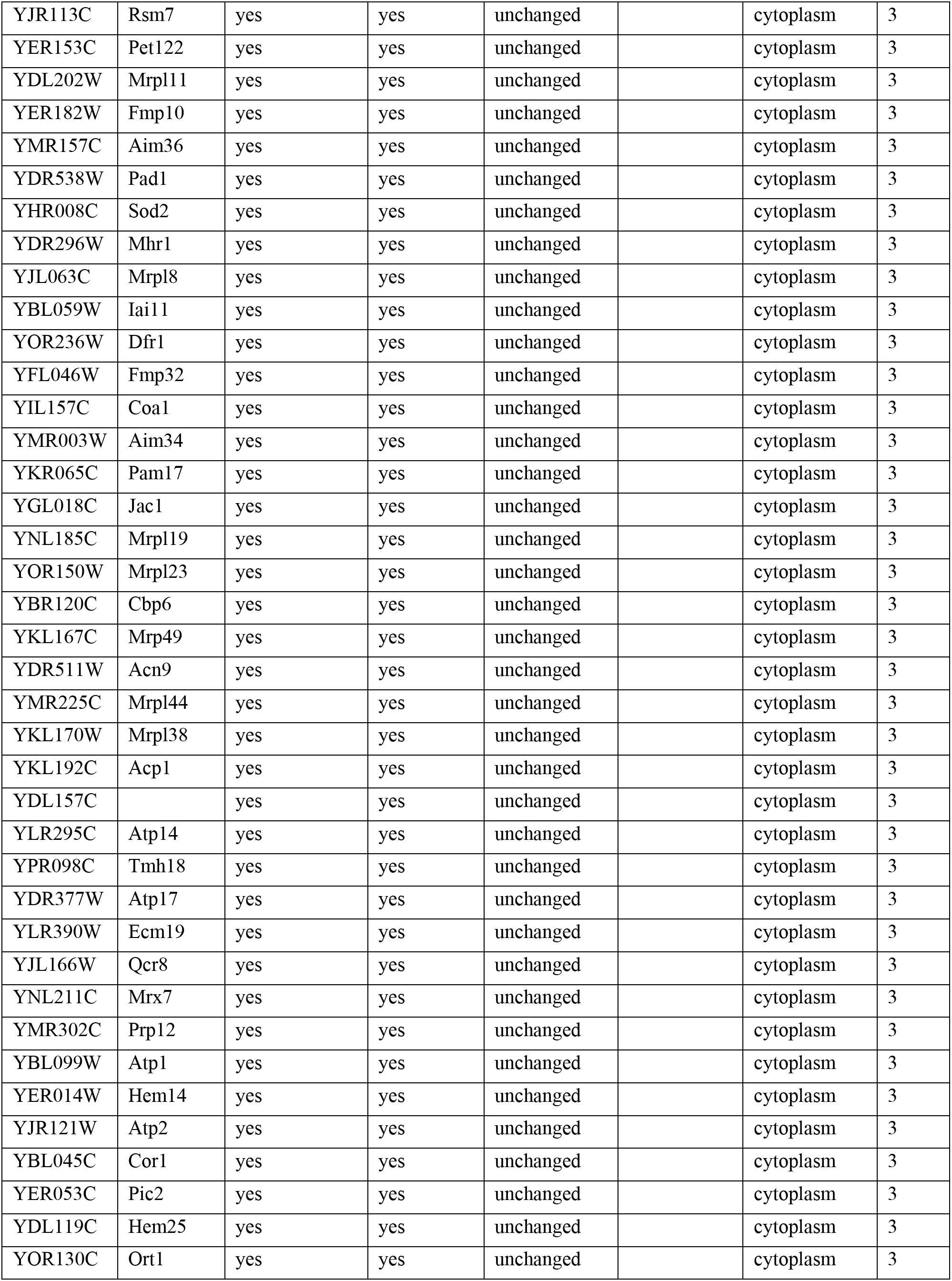
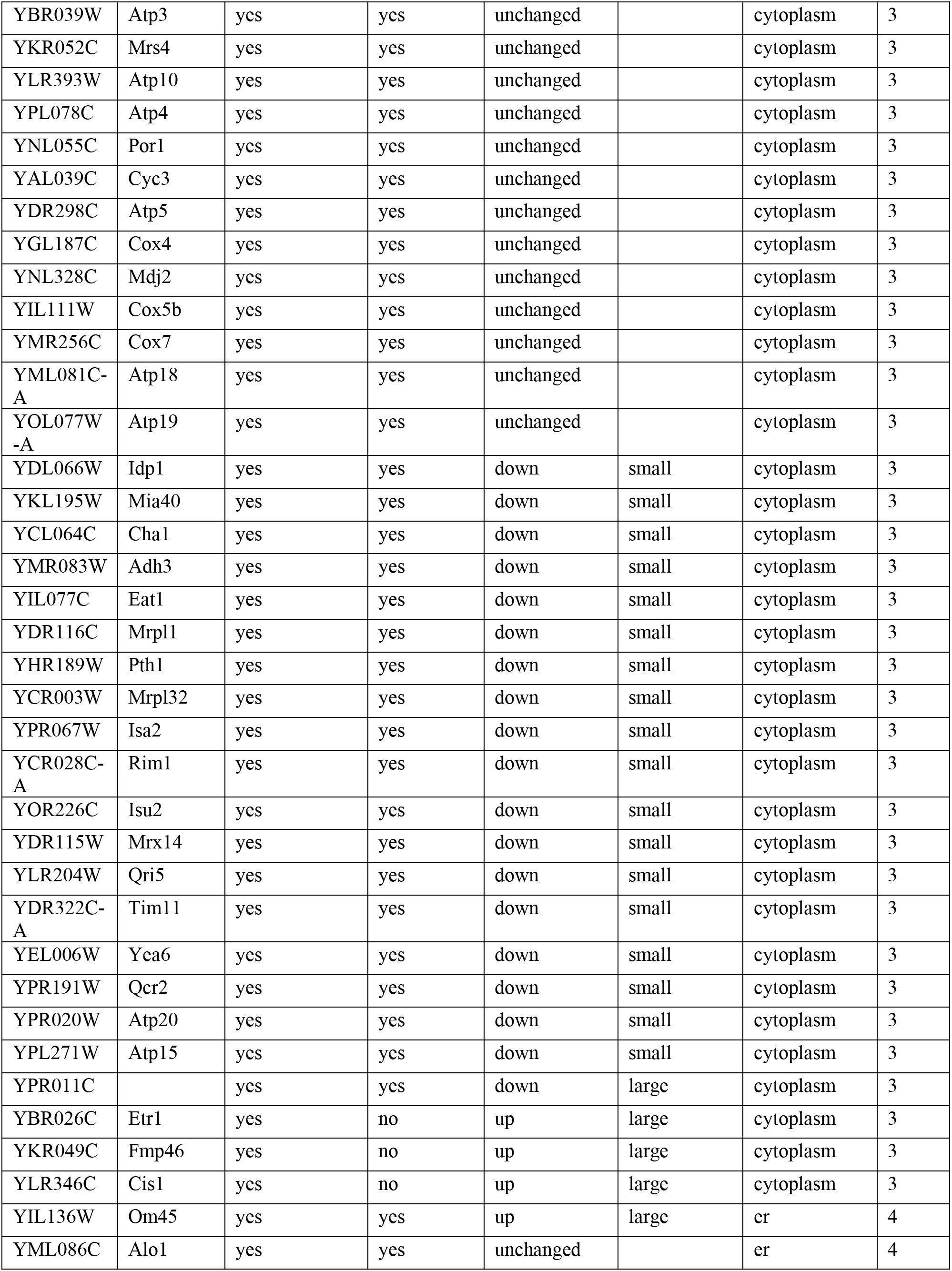
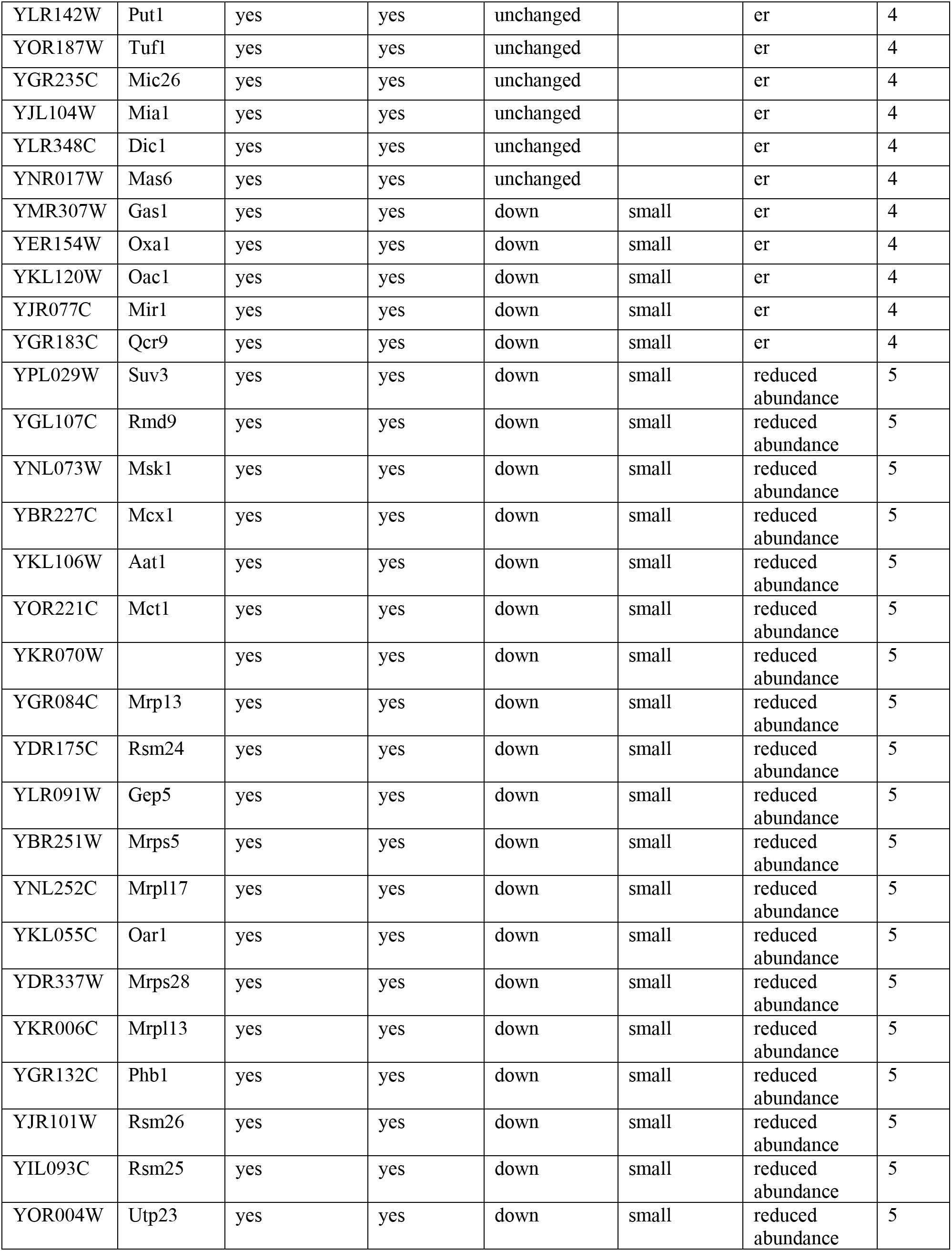
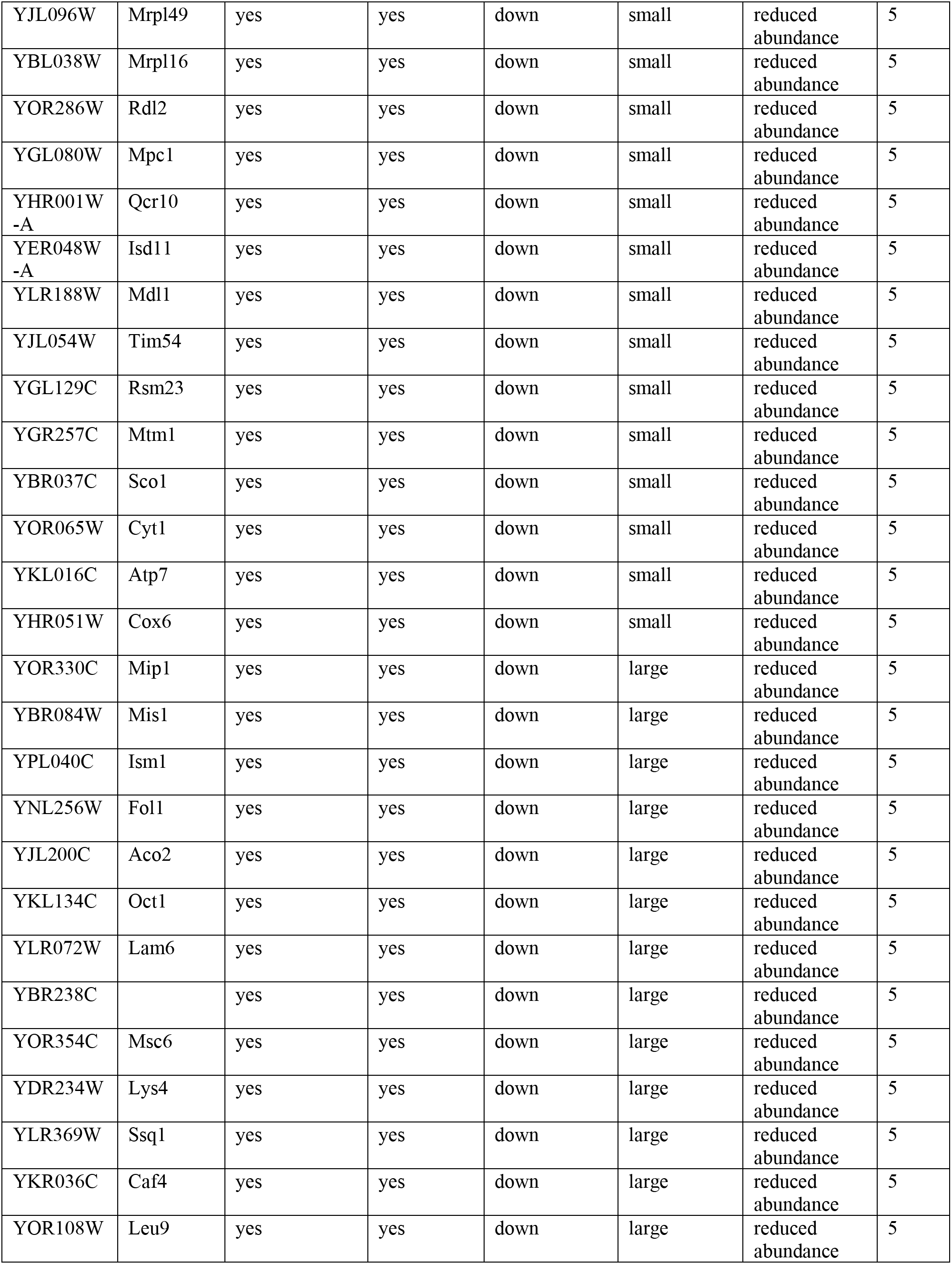
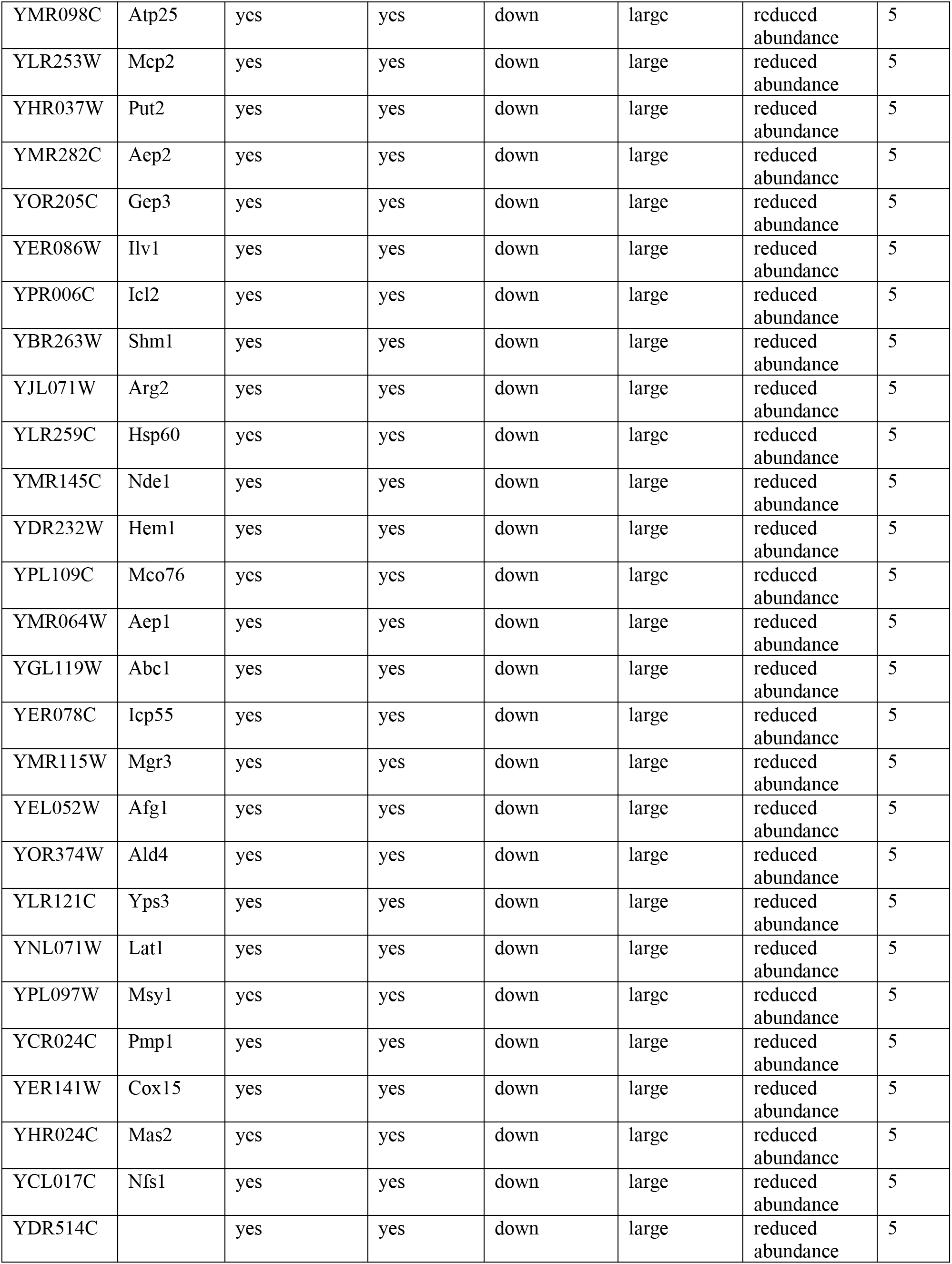
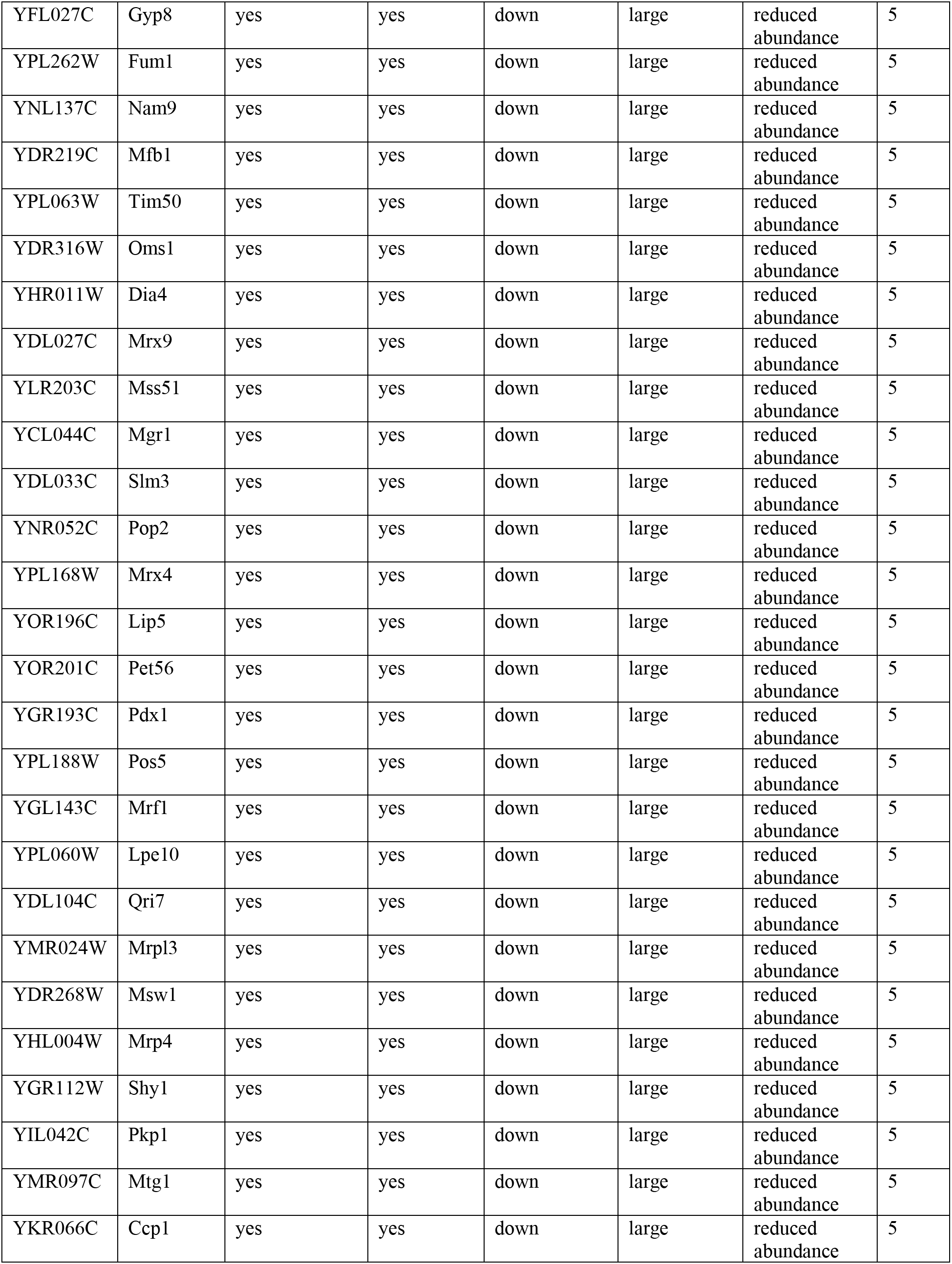
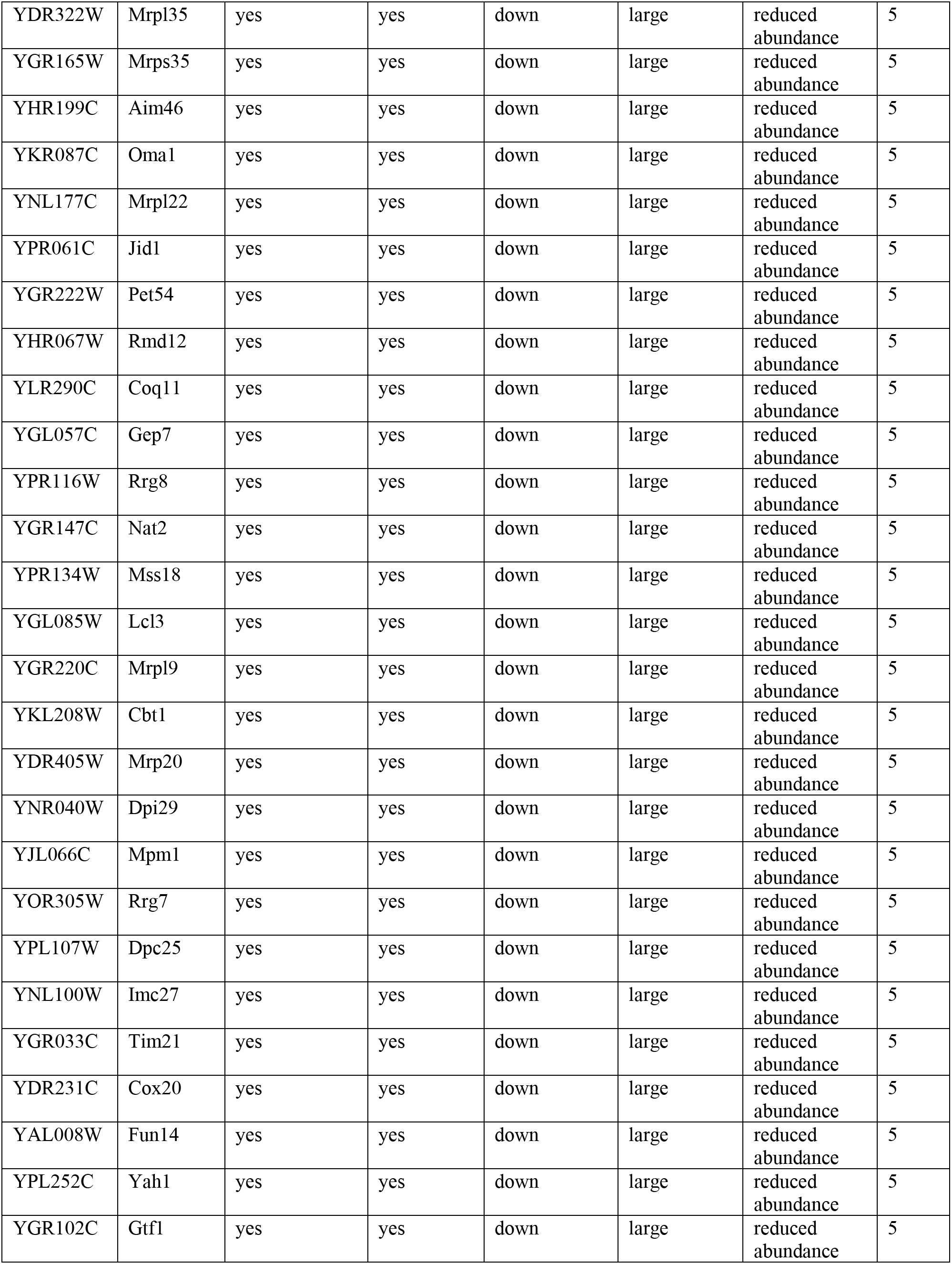
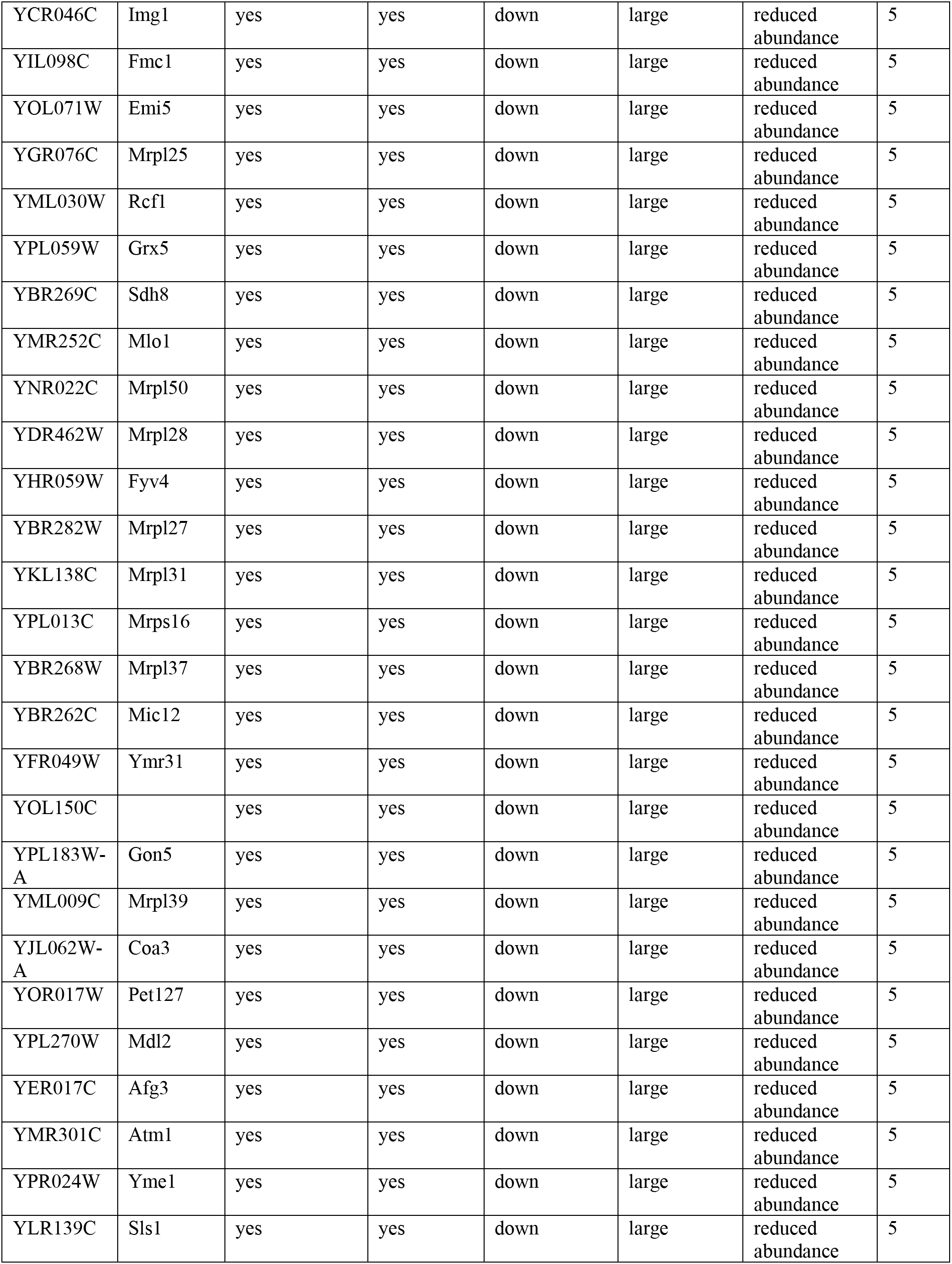
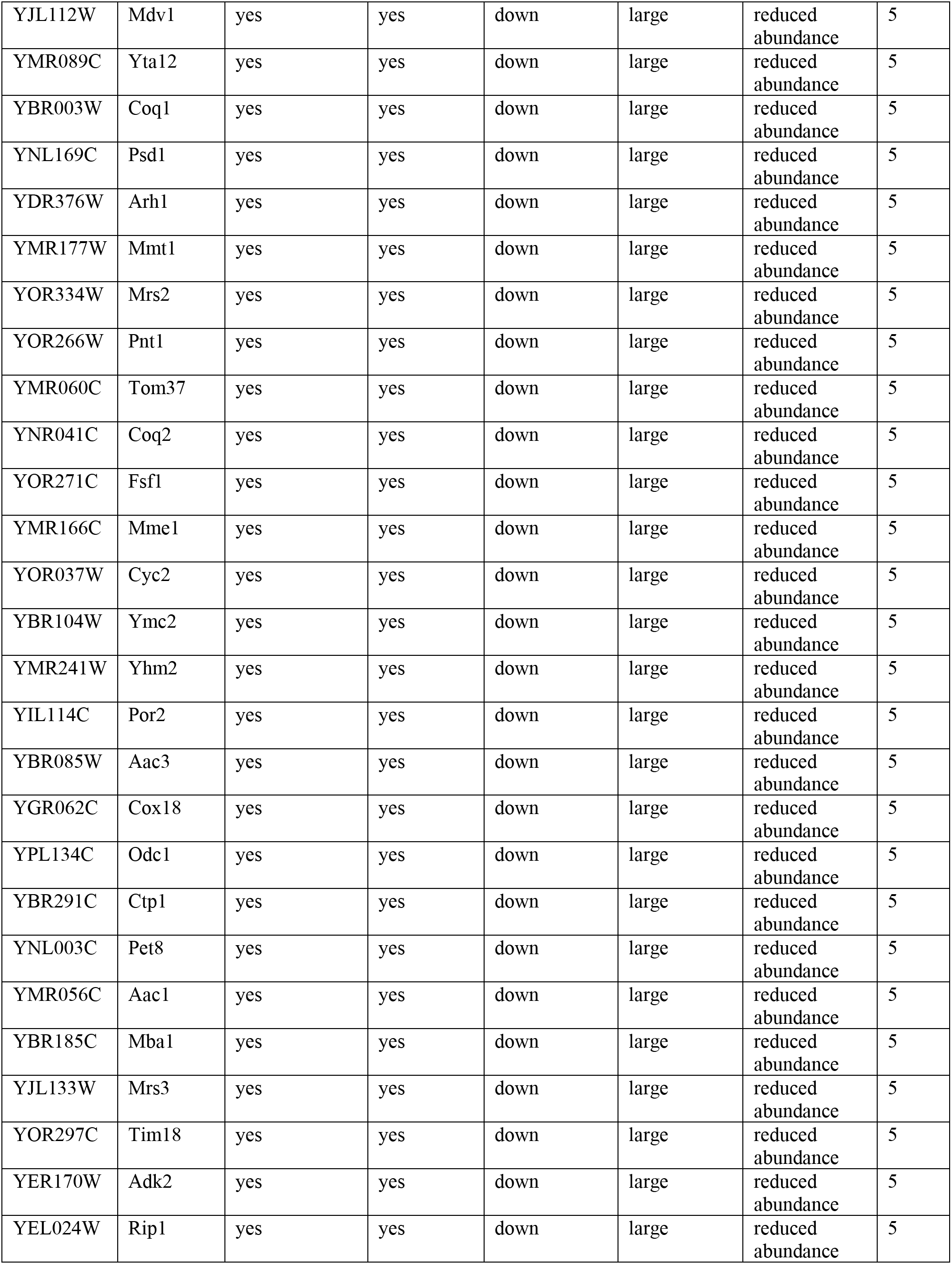
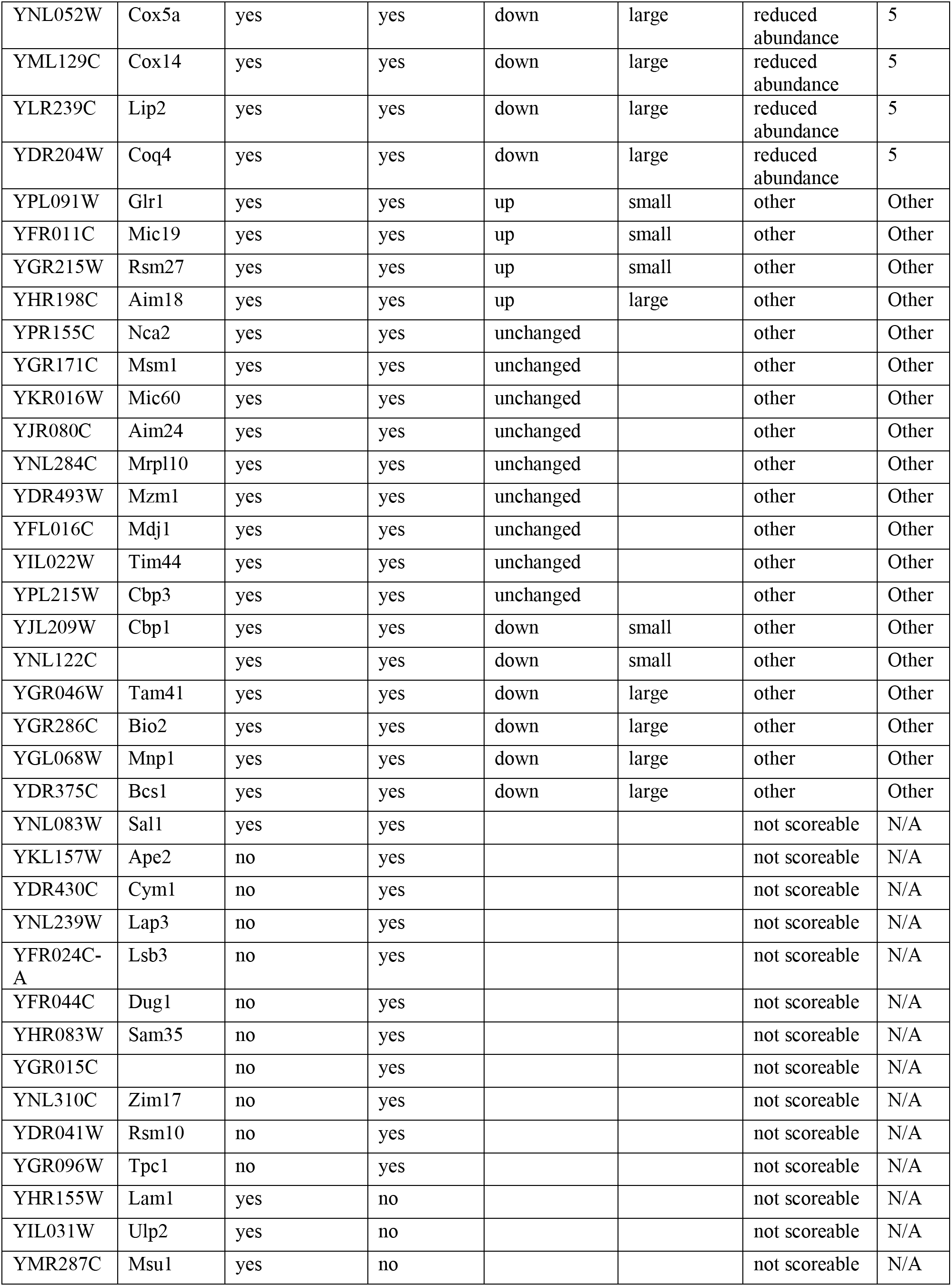
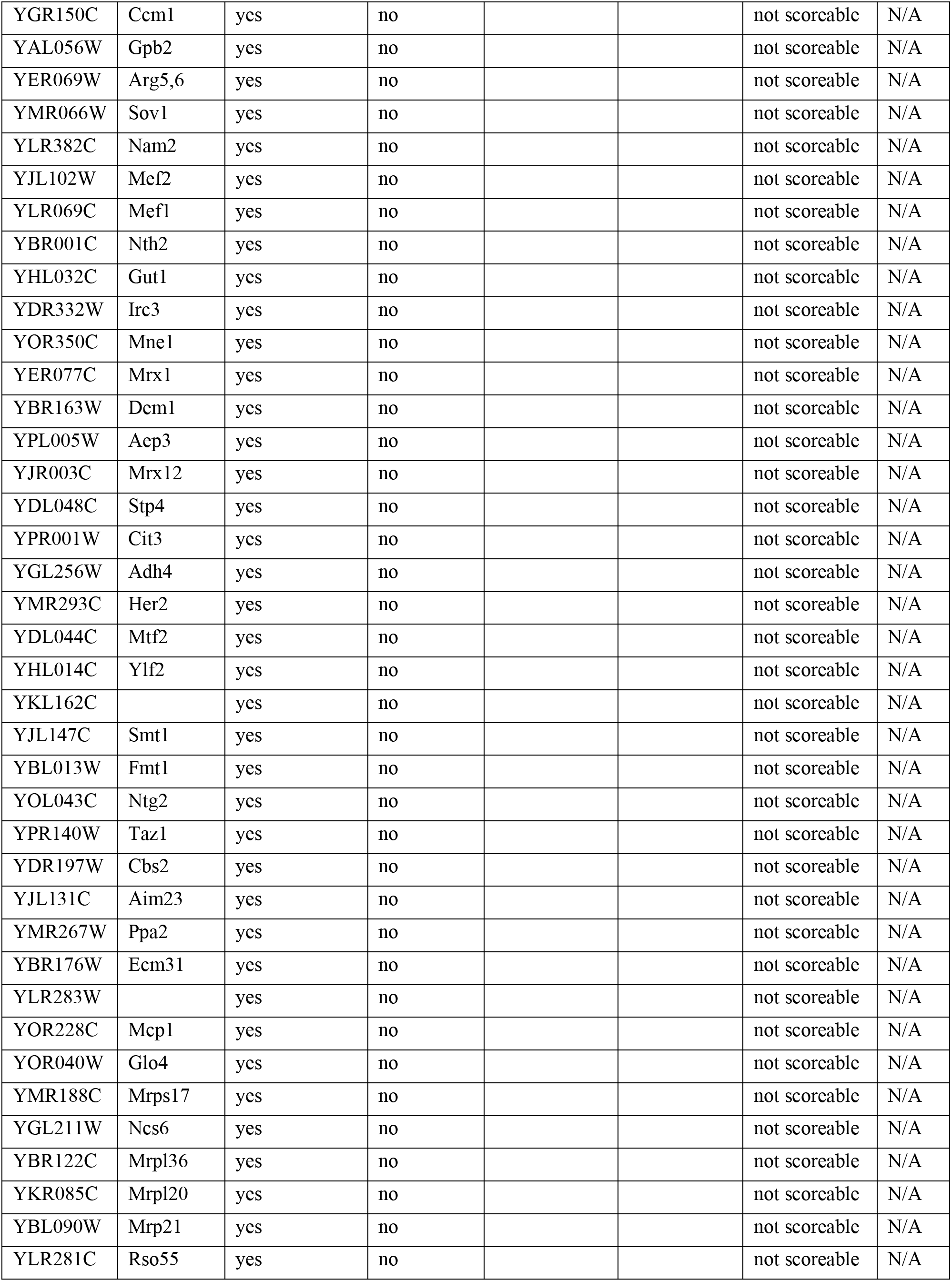
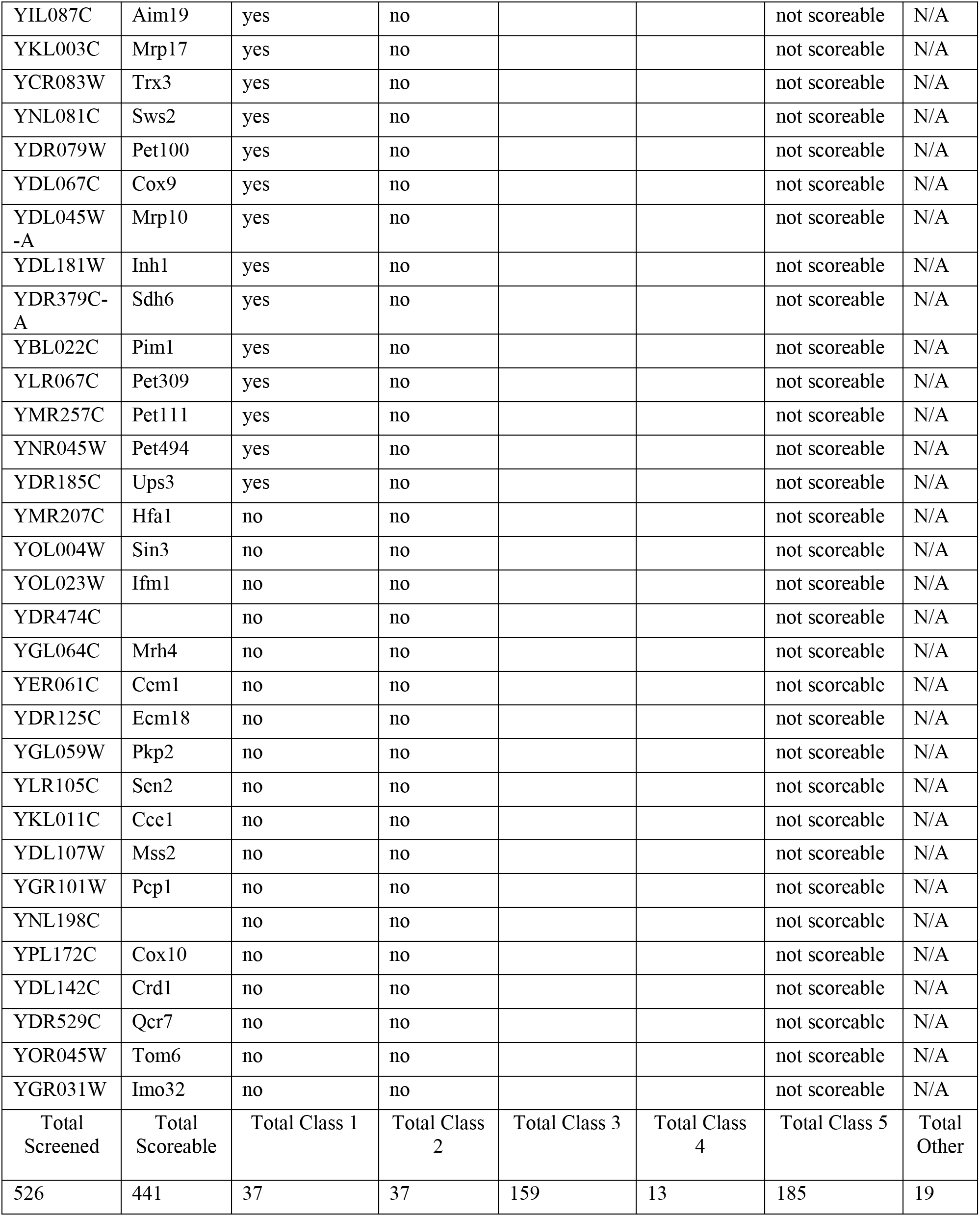
Complete list of mitochondrial protein fates upon FCCP treatment.

**Table S2.** Yeast strains used in this study.

| Strain | Genotype |
| --- | --- |
| BY4741 | MATa his3Δ leu2Δ ura3Δ met15Δ |
| AHY3354 | MATa his3Δ leu2Δ ura3Δ met15Δ TOM70-mCherry:KanMX MIR1-yeGFP:HisMX |
| AHY3742 | MATa his3Δ leu2Δ ura3Δ met15Δ TOM70-mCherry:KanMX COX15-yeGFP:HisMX |
| AHY3746 | MATa his3Δ leu2Δ ura3Δ met15Δ TOM70-mCherry:KanMX LAT1-yeGFP:HisMX |
| AHY3857 | MATa his3Δ leu2Δ ura3Δ met15Δ TOM70-mCherry:KanMX COX15-yeGFP:HisMX pdr5Δ::URA3 |
| AHY3861 | MATa his3Δ leu2Δ ura3Δ met15Δ TOM70-mCherry:KanMX LAT1-yeGFP:HisMX pdr5Δ::URA3 |
| AHY3934 | MATa/MATa his3Δ1/his3Δ1 leu2Δ0/leu2Δ0 ura3Δ0/ura3Δ0 lys2Δ0/+ met15Δ0/+ Term<sub>cyc1</sub>:URA3-P<sub>GPD/TDH3</sub>-cre-EBD78:Term<sub>cyc1</sub>/+ ILV2-V5-loxP-HA-GFP-HygX-loxP-T7-mRFP-KanMX/+ |
| AHY4042 | MATa his3Δ leu2Δ ura3Δ met15Δ ILV2-yeGFP:HisMX |
| AHY4389 | MATa his3Δ leu2Δ ura3Δ met15Δ ILV2-yeGFP:HisMX pdr5Δ::URA3 |
| AHY4628 | MATa his3Δ leu2Δ met15Δ URA3::CMV-tTA TOM70-mCherry:KanMX ILV2-yeGFP:HisMX |
| AHY4737 | MATa his3Δ leu2Δ ura3Δ met15Δ TOM70-mCherry:KanMX ACP1-yeGFP:HisMX |
| AHY4739 | MATa his3Δ leu2Δ ura3Δ met15Δ ILV2-yeGFP:HisMX TOM70-mCherry:KanMX |
| AHY4945 | MATa his3Δ leu2Δ ura3Δ met15Δ ILV2-3xHA:HisMX |
| AHY4949 | MATa his3Δ leu2Δ ura3Δ met15Δ DLD1-yeGFP:HisMX |
| AHY4951 | MATa his3Δ leu2Δ ura3Δ met15Δ DLD2-yeGFP:HisMX |
| AHY4959 | MATa his3Δ leu2Δ ura3Δ met15Δ DLD1-yeGFP:HisMX pdr5Δ::URA3 |
| AHY4961 | MATa his3Δ leu2Δ ura3Δ met15Δ DLD2-yeGFP:HisMX pdr5Δ::URA3 |
| AHY4963 | MATa his3Δ leu2Δ met15Δ URA3::CMV-tTA TOM70-mCherry:KanMX DLD1-yeGFP:HisMX |
| AHY4965 | MATa his3Δ leu2Δ met15Δ URA3::CMV-tTA TOM70-mCherry:KanMX DLD2-yeGFP:HisMX |
| AHY4971 | MATa his3Δ leu2Δ ura3Δ met15Δ ILV2-3xHA:HisMX pdr5Δ::URA3 |
| AHY5044 | MATa his3Δ leu2Δ ura3Δ met15Δ san1Δ::NatMX |
| AHY5047 | MATa his3Δ leu2Δ ura3Δ lys2Δ ubr1Δ::URA3 doa10Δ::HygMX |
| AHY5048 | MATa his3Δ leu2Δ ura3Δ met15Δ doa10Δ::HygMX |
| AHY5049 | MATa his3Δ leu2Δ ura3Δ met15Δ lys2Δ san1Δ::NatMX doa10Δ::HygMX |
| AHY5053 | MATa his3Δ leu2Δ ura3Δ lys2Δ ubr1Δ::URA3 |
| AHY5055 | MATa his3Δ leu2Δ ura3Δ met15Δ san1Δ::NatMX ubr1Δ::URA3 |
| AHY5056 | MATa his3Δ leu2Δ ura3Δ met15Δ lys2Δ san1Δ::NatMX ubr1Δ::URA3 doa10Δ::HygMX |
| AHY5058 | MATa his3Δ leu2Δ ura3Δ met15Δ lys2Δ san1Δ::NatMX ubr1Δ::URA3 doa10Δ::HygMX DLD1-yeGFP:HisMX |
| AHY5060 | MATa his3Δ leu2Δ ura3Δ met15Δ lys2Δ san1Δ::NatMX ubr1Δ::URA3 doa10Δ::HygMX DLD2-yeGFP:HisMX |
| AHY5062 | MATa his3Δ leu2Δ ura3Δ met15Δ lys2Δ san1Δ::NatMX ubr1Δ::URA3 doa10Δ::HygMX ILV2-yeGFP:HisMX |
| AHY6027 | MATa his3Δ leu2Δ ura3Δ met15Δ san1Δ::NatMX DLD1-yeGFP:HisMX |
| AHY6029 | MATa his3Δ leu2Δ ura3Δ lys2Δ ubr1Δ::URA3 DLD1-yeGFP:HisMX |
| AHY6031 | MATa his3Δ leu2Δ ura3Δ met15Δ doa10Δ::HygMX DLD1-yeGFP:HisMX |
| AHY6033 | MATa his3Δ leu2Δ ura3Δ met15Δ san1Δ::NatMX ubr1Δ::URA3 DLD1-GFP:HisMX |
| AHY6035 | MATa his3Δ leu2Δ ura3Δ met15Δ lys2Δ san1Δ::NatMX doa10Δ::HygMX DLD1-yeGFP:HisMX |
| AHY6037 | MATa his3Δ leu2Δ ura3Δ lys2Δ ubr1Δ::URA3 doa10Δ::HygMX DLD1-yeGFP:HisMX |
| AHY6039 | MATa his3Δ leu2Δ ura3Δ met15Δ san1Δ::NatMX DLD2-yeGFP:HisMX |
| AHY6041 | MATa his3Δ leu2Δ ura3Δ lys2Δ ubr1Δ::URA3 DLD2-yeGFP:HisMX |
| AHY6043 | MATa his3Δ leu2Δ ura3Δ met15Δ doa10Δ::HygMX DLD2-yeGFP:HisMX |
| AHY6045 | MATa his3Δ leu2Δ ura3Δ met15Δ san1Δ::NatMX ubr1Δ::URA3 DLD2-yeGFP:HisMX |
| AHY6047 | MATa his3Δ leu2Δ ura3Δ met15Δ lys2Δ san1Δ::NatMX doa10Δ::HygMX DLD2-yeGFP:HisMX |
| AHY6049 | MATa his3Δ leu2Δ ura3Δ lys2Δ ubr1Δ::URA3 doa10Δ::HygMX DLD2-yeGFP:HisMX |
| AHY6051 | MATa his3Δ leu2Δ ura3Δ met15Δ san1Δ::NatMX ILV2-yeGFP:HisMX |
| AHY6053 | MATa his3Δ leu2Δ ura3Δ lys2Δ ubr1Δ::URA3 ILV2-yeGFP:HisMX |
| AHY6055 | MATa his3Δ leu2Δ ura3Δ met15Δ doa10Δ::HygMX ILV2-yeGFP:HisMX |
| AHY6057 | MATa his3Δ leu2Δ ura3Δ met15Δ san1Δ::NatMX ubr1Δ::URA3 ILV2-yeGFP:HisMX |
| AHY6059 | MATa his3Δ leu2Δ ura3Δ met15Δ lys2Δ san1Δ::NatMX doa10Δ::HygMX ILV2-yeGFP:HisMX |
| AHY6061 | MATa his3Δ leu2Δ ura3Δ lys2Δ ubr1Δ::URA3 doa10Δ::HygMX ILV2-yeGFP:HisMX |
| AHY6063 | MATa his3Δ leu2Δ ura3Δ met15Δ lys2Δ san1Δ::NatMX ubr1Δ::URA3 doa10Δ::HygMX LAT1-yeGFP:HisMX |
| AHY6408 | MATa his3Δ leu2Δ ura3Δ met15Δ lys2Δ san1Δ::NatMX ubr1Δ::URA3 doa10Δ::HygMX ΔPDR5::G418, ILV2-3xHA:HixMX |
| AHY6802 | MATa his3Δ leu2Δ met15Δ P<sub>TOM40</sub>::NatMX-tet07-TATA URA3::CMV-tTA TOM70-mCherry:KanMX ILV2-yeGFP:HisMX |
| AHY6804 | MATa his3Δ leu2Δ met15Δ P<sub>TOM40</sub>::NatMX-tet07-TATA URA3::CMV-tTA TOM70-mCherry:KanMX DLD1-yeGFP:HisMX |
| AHY6806 | MATa his3Δ leu2Δ met15Δ P<sub>TOM40</sub>::NatMX-tet07-TATA URA3::CMV-tTA TOM70-mCherry:KanMX DLD2-GFP:HisMX |
| AHY6808 | MATa his3Δ leu2Δ met15Δ P<sub>TOM40</sub>::NatMX-tet07-TATA URA3::CMV-tTA TOM70-mCherry:KanMX MIR1-yeGFP:HisMX |
| AHY6864 | MATa his3Δ leu2Δ met15Δ P<sub>TOM40</sub>::NatMX-tet07-TATA URA3::CMV-tTA TOM70-mCherry:KanMX TOM20-yeGFP:HisMX |
| AHY6867 | MATa his3Δ leu2Δ met15Δ P<sub>TOM40</sub>::NatMX-tet07-TATA URA3::CMV-tTA TOM70-mCherry:KanMX COX15-yeGFP:HisMX |
| AHY6870 | MATa his3Δ leu2Δ met15Δ P<sub>TOM40</sub>::NatMX-tet07-TATA URA3::CMV-tTA TOM70-mCherry:KanMX ACP1-yeGFP:HisMX |
| AHY6948 | MATa his3Δ leu2Δ ura3Δ met15Δ SEC61-mCherry:KanMX MIR1-yeGFP:HisMX |
| AHY7181 | MATa his3Δ leu2Δ met15Δ URA3::CMV-tTA TOM70-mCherry:KanMX ACP1-yeGFP:HisMX |
| AHY7183 | MATa his3Δ leu2Δ met15Δ URA3::CMV-tTA TOM70-mCherry:KanMX MIR1-yeGFP:HisMX |
| AHY7187 | MATa his3Δ leu2Δ met15Δ URA3::CMV-tTA TOM70-mCherry:KanMX COX15-yeGFP:HisMX |
| AHY7226 | MATa his3Δ leu2Δ ura3Δ met15Δ TOM70-mCherry:KanMX pRS413-pGPD-COX15-GFP |
| AHY7228 | MATa his3Δ leu2Δ ura3Δ met15Δ TOM70-mCherry:KanMX pRS413-GPD-ΔMTS (ΔN1-65) COX15-GFP |
| AHY7582 | MATa his3Δ leu2Δ ura3Δ met15Δ TOM70-mCherry:KanMX TOM20-yeGFP:HisMX |
| AHY7584 | MATa his3Δ leu2Δ ura3Δ met15Δ TOM70-mCherry:KanMX DLD1-yeGFP:HisMX |
| AHY7586 | MATa his3Δ leu2Δ ura3Δ met15Δ TOM70-mCherry:KanMX DLD2-yeGFP:HisMX |
| AHY7594 | MATa his3Δ leu2Δ ura3Δ met15Δ lys2Δ san1Δ::NatMX ubr1Δ::URA3 doa10Δ::HygMX DLD1-yeGFP:HisMX TOM70-mCherry:KanMX |
| AHY7596 | MATa his3Δ leu2Δ ura3Δ met15Δ lys2Δ san1Δ::NatMX ubr1Δ::URA3 doa10Δ::HygMX DLD2-yeGFP:HisMX TOM70-mCherry:KanMX |
| AHY7598 | MATa his3Δ leu2Δ ura3Δ met15Δ lys2Δ san1Δ::NatMX, ubr1Δ::URA3 doa10Δ::HygMX ILV2-yeGFP:HisMX TOM70-mCherry:KanMX |
| AHY7742 | MATa his3Δ leu2Δ met15Δ URA3::CMV-tTA TOM70-mCherry:KanMX TOM20-yeGFP:HisMX |
| AHY7875 | MATa his3Δ leu2Δ ura3Δ met15Δ TOM70-mCherry:KanMX pRS413-GPD-ΔMTS (ΔN1-55)ILV2-GFP |
| AHY7876 | MATa his3Δ leu2Δ ura3Δ met15Δ TOM70-mCherry:KanMX pRS413-GPD-MTS<sub>ILV2</sub>-GFP |
| AHY7965 | MATa his3Δ leu2Δ ura3Δ met15Δ TOM70-mCherry:KanMX pRS413-pGPD-MTS<sub>COX15</sub>-GFP |
| AHY7967 | MATa his3Δ leu2Δ ura3Δ met15Δ TOM70-mCherry:KanMX pRS413-GPD-ΔMTS (ΔN 1-28) LAT1-GFP |
| AHY7969 | MATa his3Δ leu2Δ ura3Δ met15Δ TOM70-mCherry:KanMX pRS413-pGPD-LAT1-G |
| AHY8001 | MATa his3Δ leu2Δ ura3Δ met15Δ lys2Δ san1Δ::NatMX, ubr1Δ::URA3 doa10Δ::HygMX TOM70-mCherry:KanMX pRS413-GPD-ΔMTS (ΔN1-55)ILV2-GFP |
| AHY8003 | MATa his3Δ leu2Δ ura3Δ met15Δ lys2Δ san1Δ::NatMX ubr1Δ::URA3 doa10Δ::HygMX TOM70-mCherry:KanMX pRS413-pGPD-MTS<sub>ILV2</sub>-GFP |
| AHY8008 | MATa his3Δ leu2Δ ura3Δ met15Δ TOM70-mCherry:KanMX pRS413-pGPD-MTS<sub>LAT1</sub>-GFP |
| AHY8027 | MATa his3Δ leu2Δ ura3Δ met15Δ TOM70-mCherry:KanMX pRS413-pGPD-ILV2-GFP |
| AHY8031 | MATa his3Δ leu2Δ ura3Δ met15Δ lys2Δ san1Δ::NatMX ubr1Δ::URA3 doa10Δ::HygMX TOM70-mCherry:KanMX pRS413-pGPD-ILV2-GFP |
| AHY8043 | MATa his3Δ leu2Δ ura3Δ met15Δ lys2Δ san1Δ::NatMX ubr1Δ::URA3 doa10Δ::HygMX Tom70-mCherry:KanMX pRS413-pGPD-DLD2-GFP |
| AHY8345 | MATa his3Δ leu2Δ ura3Δ met15Δ TOM70-GFP:KanMX TIM50-mCherry:KanMX ilv2Δ::URA3 |
| AHY8557 | MATa his3Δ leu2Δ ura3Δ met15Δ TOM70-mCherry:KanMX pRS413-GPD-ΔMTS (ΔN1-35) DLD2-GFP |
| AHY8559 | MATa his3Δ leu2Δ ura3Δ met15Δ lys2Δ san1Δ::NatMX, ubr1Δ::URA3 doa10Δ::HygMX TOM70-mCherry:KanMX pRS413-GPD-ΔMTS (ΔN1-35) DLD2-GFP |
| AHY8561 | MATa his3Δ leu2Δ ura3Δ met15Δ TOM70-mCherry-KanMX pRS413-pGPD-MTS<sub>DLD2</sub>-GFP |
| AHY8563 | MATa his3Δ leu2Δ ura3Δ met15Δ lys2Δ san1Δ::NatMX ubr1Δ::URA3 doa10Δ::HygMX TOM70-mCherry:KanMX pRS413-pGPD-MTS<sub>DLD2</sub>-GFP |
| AHY8671 | MATa his3Δ leu2Δ ura3Δ met15Δ TOM70-mCherry:KanMX pRS413-pGPD-DLD2-GFP |
| AHY10107 | MATa his3Δ leu2Δ ura3Δ met15Δ lys2ΔTOM70-mCherry KanMX pdr5Δ::URA3 LAT1-3XHA-HisMX |
| AHY10198 | MATa his3Δ leu2Δ ura3Δ met15Δ ILV2-3xHA:KanMX pdr5Δ::Ura3 NUP49-yeGFP:HisMX |
| AHY10267 | MATa his3Δ leu2Δ ura3Δ met15Δ ACP1-3xHA:HisMX |
| AHY10269 | MATa his3Δ leu2Δ ura3Δ met15Δ COX15-3xHA:HisMX |
| AHY10369 | MATa his3Δ leu2Δ ura3Δ met15Δ ILV2-5FLAG:KanMX TOM70-mCherry:HygMX |
| AHY10371 | MATa his3Δ leu2Δ ura3Δ met15Δ TOM20-5FLAG:KanMX TOM70-mCherry:HygMX |
| AHY10373 | MATa his3Δ leu2Δ ura3Δ met15Δ MIR1-5FLAG:KanMX TOM70-mCherry:HygMX |
| AHY10375 | MATa his3Δ leu2Δ ura3Δ met15Δ ACP1-5FLAG:KanMX TOM70-mCherry:HygMX |
| AHY10377 | MATa his3Δ leu2Δ ura3Δ met15Δ COX15-5FLAG:KanMX TOM70-mCherry:HygMX |
| AHY10381 | MATa his3Δ leu2Δ ura3Δ met15Δ MIR1-3xHA:HisMX |
| AHY10385 | MATa his3Δ leu2Δ ura3Δ met15Δ TOM20-3xHA:HisMX TOM70-mCherry:KanMX |
| AHY10437 | MATa his3Δ leu2Δ ura3Δ met15Δ COX15-3xHA:HisMX pdr5Δ::URA3 |

**Table S3.**
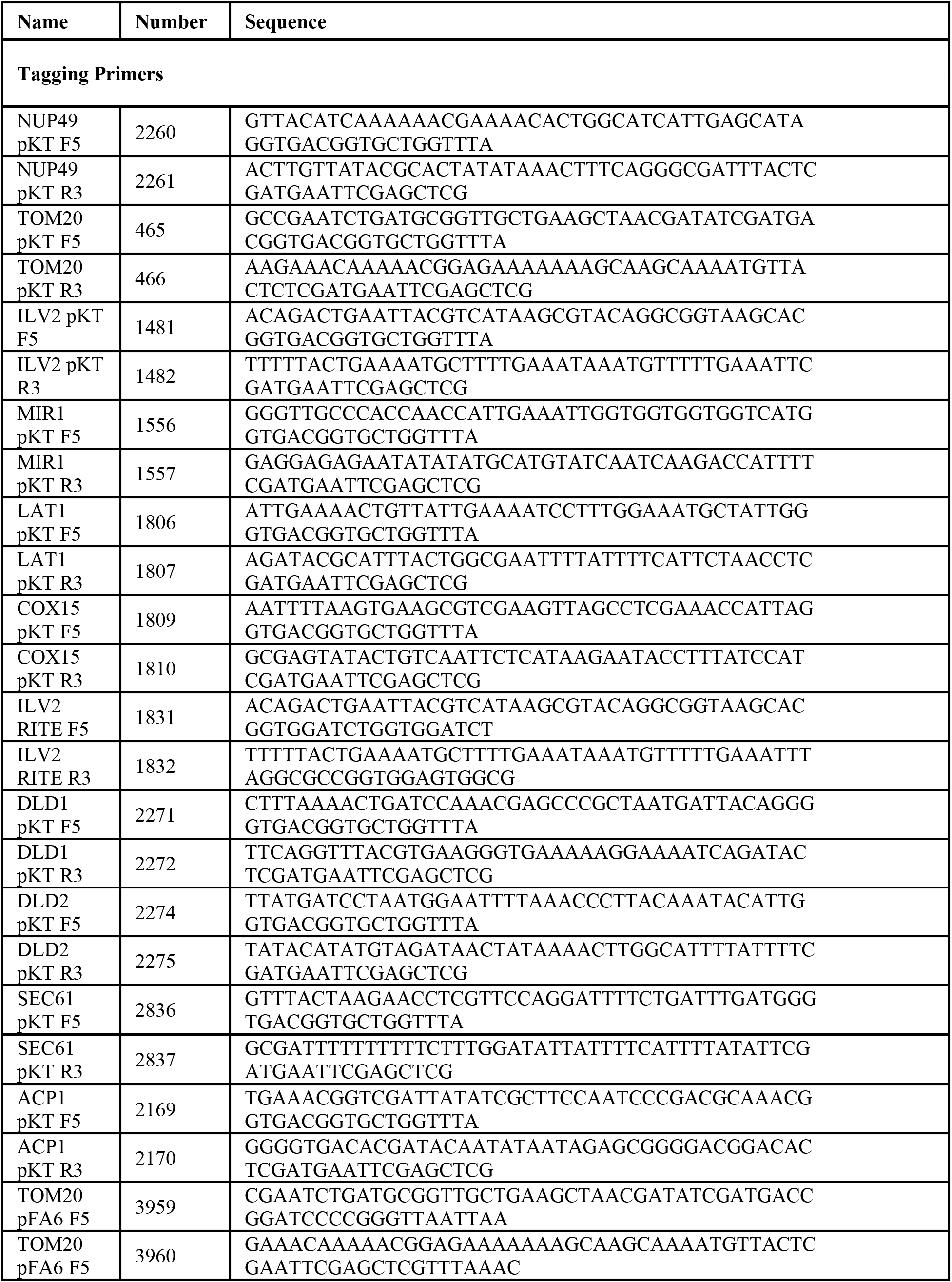
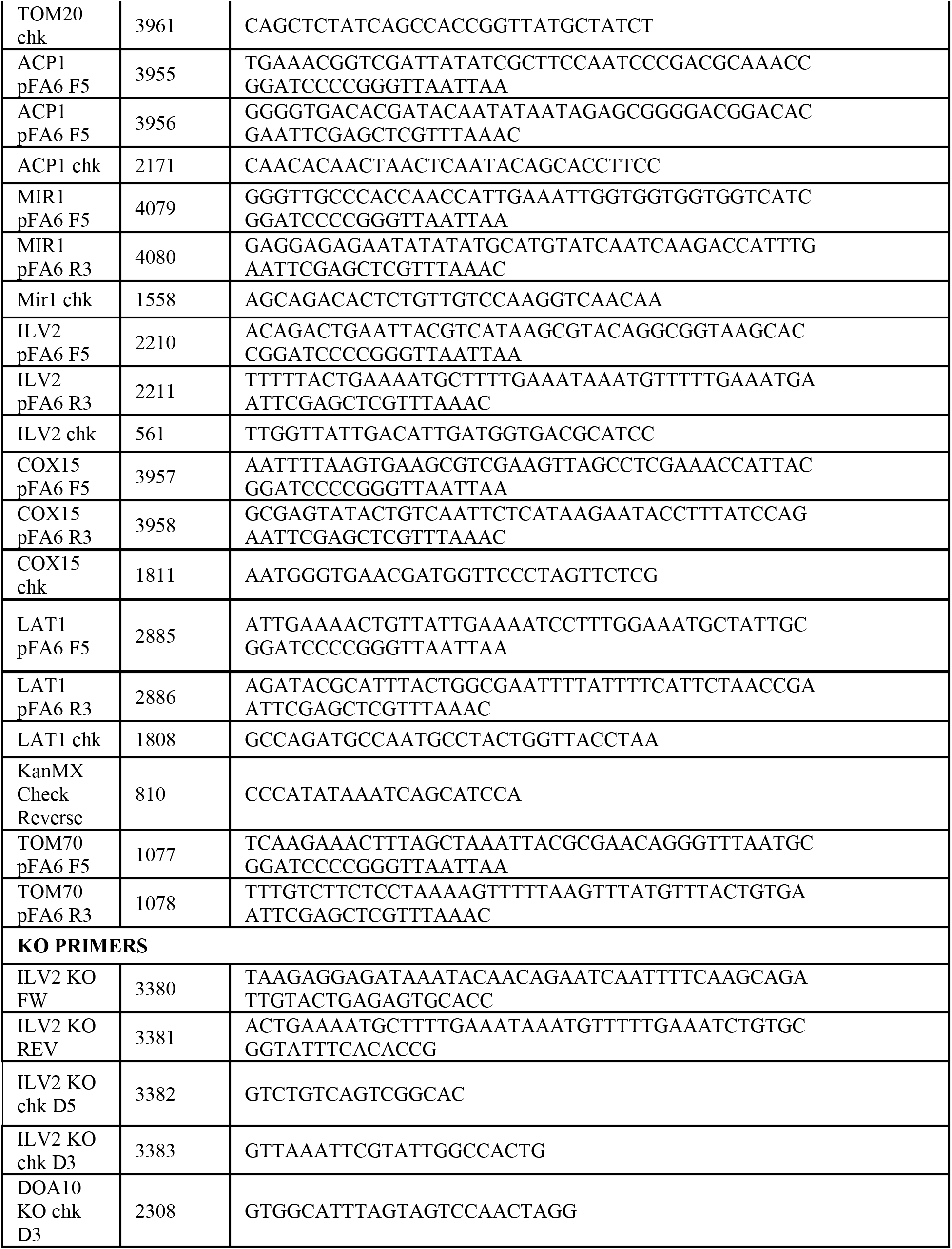
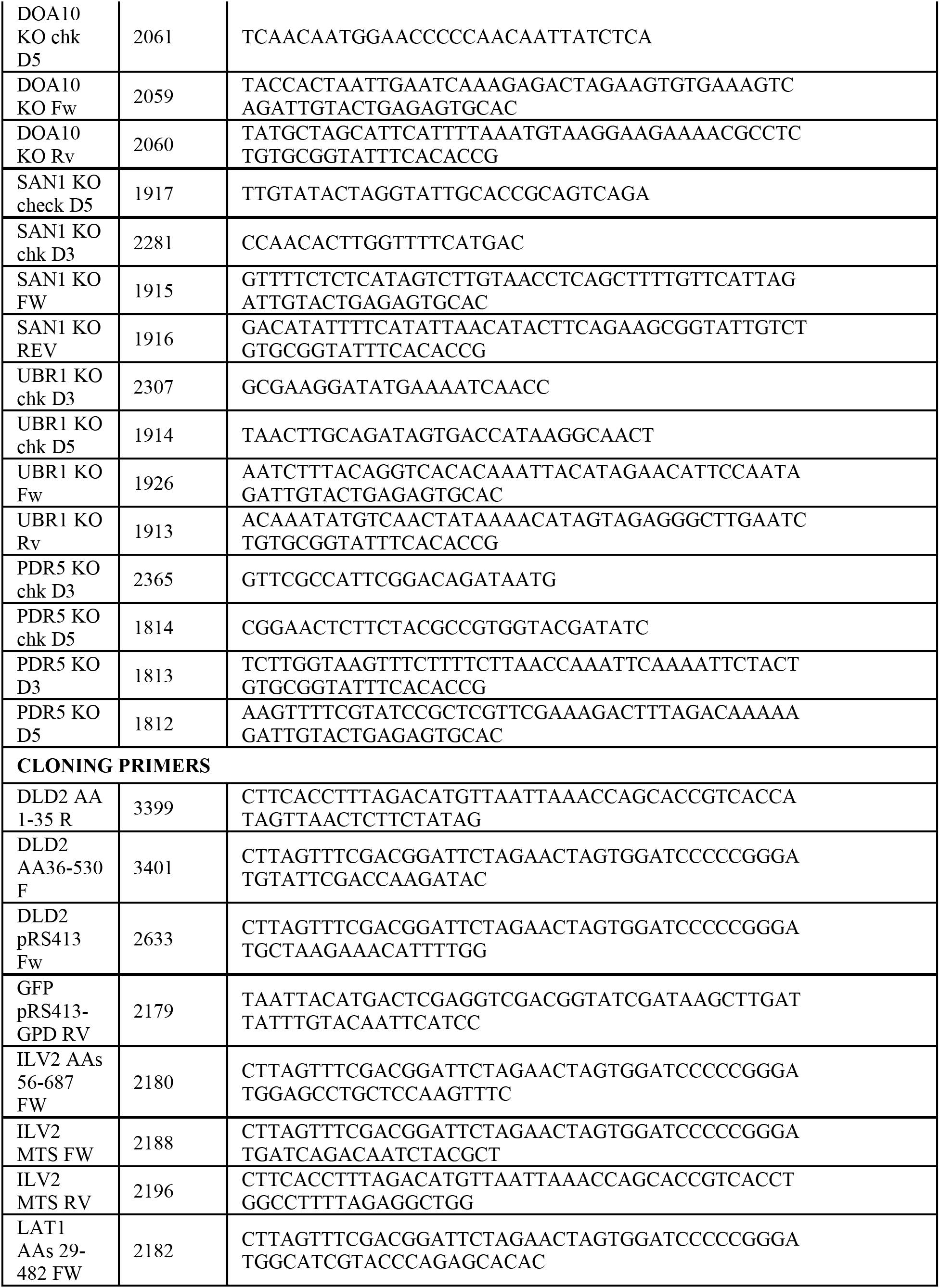
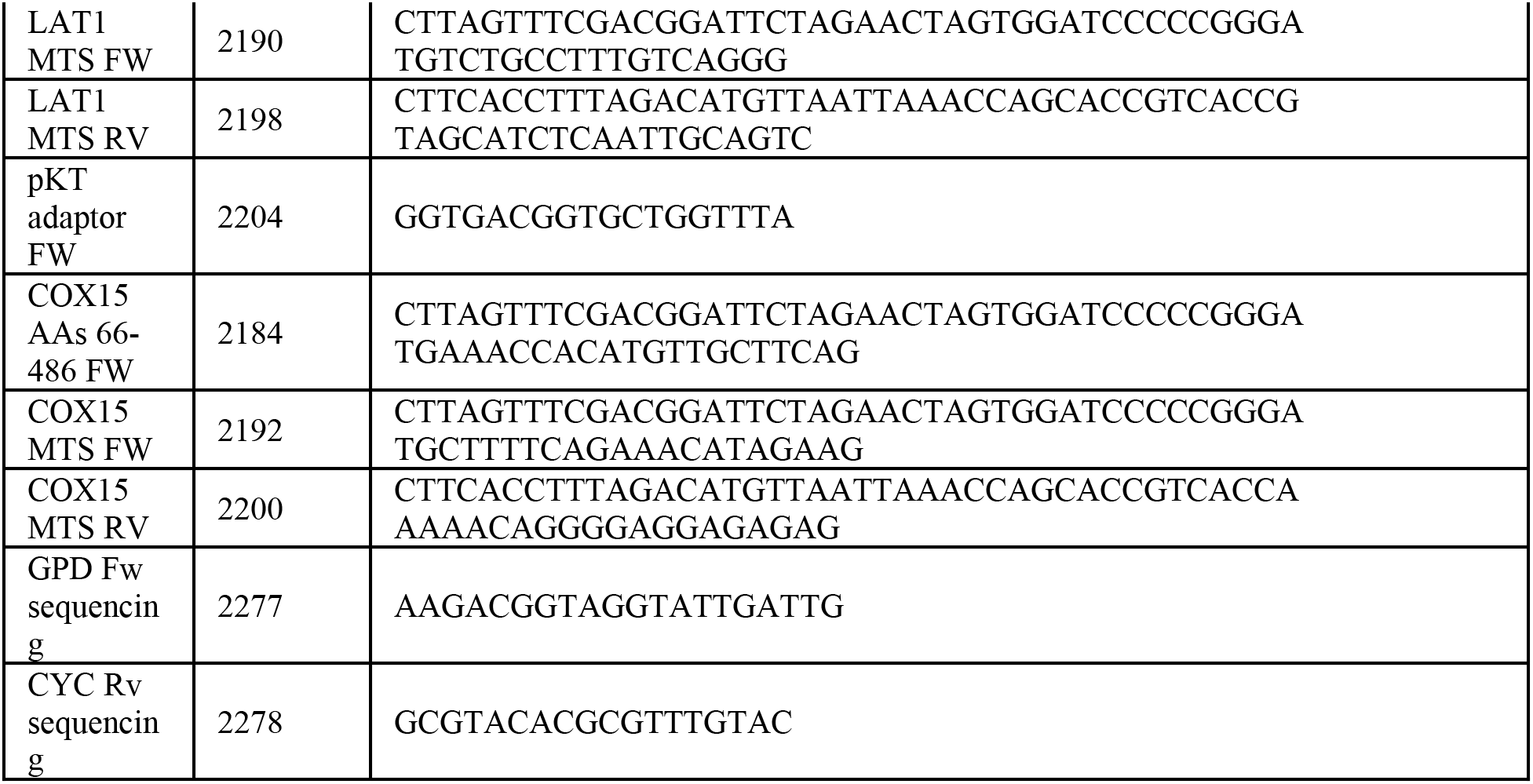
Oligos used in this study.

**Table S4.** Plasmids used in this study.

| Plasmid | Construction |
| --- | --- |
| pRS413-GPD-ILV2-GFP | pRS413-GPD cut w/ EcoRI + PCR product 1 (AHY4042 gDNA amplified w/ 2188/2179) |
| pRS413-GPD-DLD2-GFP | pRS413-GPD cut w/ EcoRI + PCR product 1 (AHY4951 gDNA amplified w/ 2633/2179) |
| pRS413-GPD-COX15-GFP | pRS413-GPD cut w/ EcoRI + PCR product 1 (AHY3742 gDNA amplified w/ 2192/2179) |
| pRS413-GPD-LAT1-GFP | pRS413-GPD cut w/ EcoRI + PCR product 1 (AHY3746 gDNA amplified w/ 2190/2179) |
| pRS413-GPD-NΔ55ILV2-GFP | pRS413-GPD cut w/ EcoRI + PCR product 1 (AHY4042 gDNA amplified w/ 2180/2179) |
| pRS413-GPD-NΔ35DLD2-GFP | pRS413-GPD cut w/ EcoRI + PCR product 1 (pRS413-GPD-DLD2-GFP amplified w/ 3401/2179) |
| pRS413-GPD-NΔ65COX15-GFP | pRS413-GPD cut w/ EcoRI + PCR product 1 (AHY3742 gDNA amplified w/ 2184/2179) |
| pRS413-GPD-NΔ28LAT1-GFP | pRS413-GPD cut w/ EcoRI + PCR product 1 (AHY3746 gDNA amplified w/ 2182/2179) |
| pRS413-GPD-MTS<sub>ILV2</sub>-GFP | pRS413-GPD cut w/ EcoRI + PCR product 1 (BY4741 gDNA amplified w/ 2188/2196) + PCR product 2 (pKT128 amplified w/ 2204/2179) |
| pRS413-GPD-MTS<sub>DLD2</sub>-GFP | pRS413-GPD cut w/ EcoRI + PCR product 1 (pRS413-GPD-DLD2-GFP amplified w/ 2633/3399) + PCR product 2 (pKT128 amplified w/ 2204/2179) |
| pRS413-GPD-MTS<sub>COX15</sub>-GFP | pRS413-GPD cut w/ EcoRI + PCR product 1 (BY4741 gDNA amplified w/ 2192/2200) + PCR product 2 (pKT128 amplified w/ 2204/2179) |
| pRS413-GPD-MTS<sub>LAT1</sub>-GFP | pRS413-GPD cut w/ EcoRI + PCR product 1 (BY4741 gDNA amplified w/ 2190/2198) + PCR product 2 (pKT128 amplified w/ 2204/2179) |

## Notes

### Competing Interest Statement

The authors have declared no competing interest.

